# Designing microplate layouts using artificial intelligence

**DOI:** 10.1101/2022.03.31.486595

**Authors:** María Andreína Francisco Rodríguez, Jordi Carreras Puigvert, Ola Spjuth

## Abstract

Microplates are indispensable in large-scale biomedical experiments but the physical location of samples and controls on the microplate can significantly affect the resulting data and quality metric values. We introduce a new method based on constraint programming for designing microplate layouts that reduces unwanted bias and limits the impact of batch effects after error correction and normalisation. We demonstrate that our method applied to dose-response experiments leads to more accurate regression curves and lower errors when estimating IC_50_/EC_50_, and for drug screening leads to increased sensitivity, when compared to random layouts. It also reduces the risk of inflated scores from common microplate quality assessment metrics such as Z’ factor and SSMD. We make our method available via a suite of tools (PLAID) including a reference constraint model, a web application, and Python notebooks to evaluate and compare designs when planning microplate experiments.

## Main

In the era of data-driven life science, the amounts of data produced are continuously expanding, and artificial intelligence techniques such as machine learning algorithms are seeing adoption for many applications in order to convert the data into actionable insights [1–5]. While in many applications the primary focus has been to obtain as much data as possible, the importance of having data of high quality cannot be understated [6–8]. For large-scale biomedical experiments, many issues related to data quality pertaining to human operations can be effectively reduced or eliminated by using automated setups and robotised equipment [9]. However, several artefacts due to physical, biological, and temporal conditions still remain, and efforts generating large quantities of data can be fruitless if in the end conclusions cannot be drawn due to data-quality issues. A common approach to increase the confidence in the data is to perform multiple technical and biological replicates, but this is associated with higher costs and longer experiments, and often leads to a trade-off between the number of samples analysed and the number of replicates per sample. Another approach is to *improve the experimental design*, with the aim to carry out the experiment in such a way that it maximises the conclusions that can be drawn from the resulting data [10].

Microplates, or microwell plates, are standard components in many biomedical experiments. They are flat plates with multiple wells used as small test tubes, organised in a 2:3 matrix. They come in a standard physical dimension to ensure compatibility with different lab equipment, and typically contain 24, 96, 384, or 1536 wells. Experiments carried out using microplates commonly exhibit plate effects [11], also known as positional effects, which are systematic variations across the geometry of a microplate (within-plate effects) or across different plates (between-plate effects) due to factors such as well location, temperature and humidity being unequally distributed, and can affect the results to the point of rendering the experiment unusable. Other factors that can contribute to experimental variation are the lab equipment, such as imprecise manual pipetting, and inconsistent or malfunctioning liquid handling instruments. Common patterns of within-plate effects include: (i) linear row effects; (ii) linear column effects; (iii) linear row and column effects; and (iv) bowl-shaped spatial effects [11]; examples are visualised in Figure 1. Identifying and correcting for both within- and between-plate effects is important in order to adjust the data so that the impact of the errors can be reduced or avoided. Various normalisation techniques have been developed to this end [12, 13], but an appropriate microplate layout is of particular importance for the normalisation to be effective [12, 14]. A *control* is a sample that has been subjected to a known treatment with the goal of accounting for the effects of variables other than what is being tested, thus increasing the reliability of the results. In particular, a *negative control* is a sample that has been subjected to a treatment that induces no effect, while a *positive control* is a sample that has been subjected to a treatment with an expected maximal response [15]. In order to mitigate plate effects and gain the most out of using control samples and error correction methods, scientists have been advocating for the use of randomised plate layouts [16, 17].

**Figure 1:**
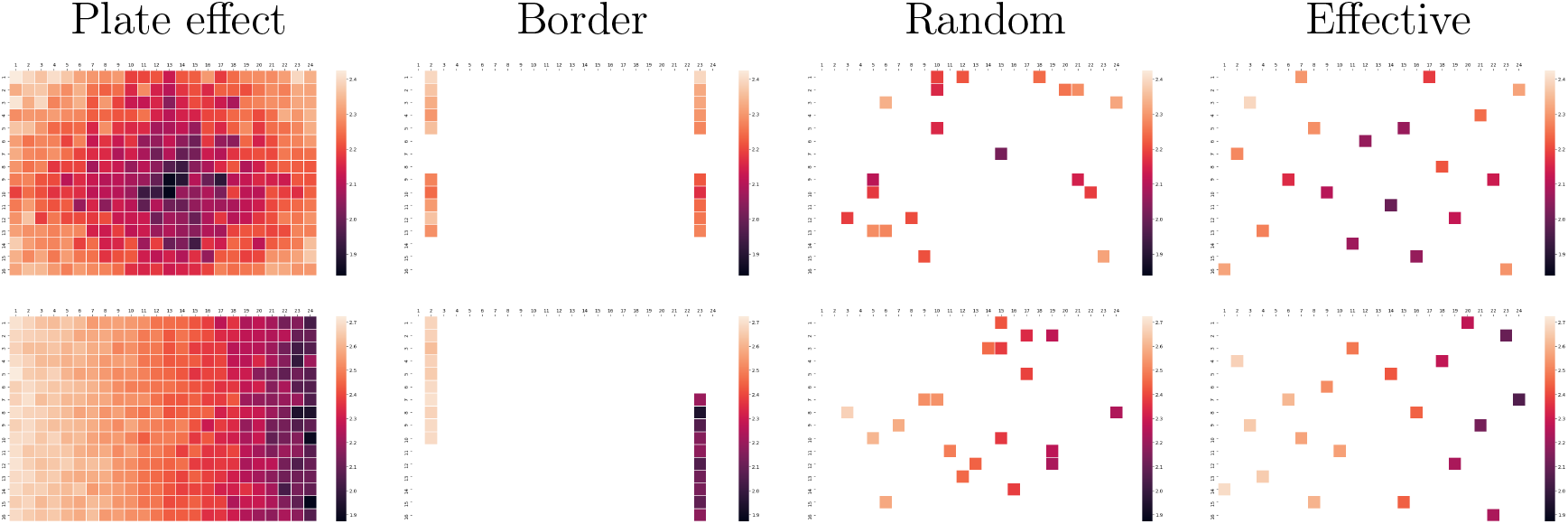
Visualisation of systematic plate effects and distributions of 20 negative controls using border, random and effective layouts. The colors indicate the intensity measured at each well. Top row: Data with strong systematic errors having a bowl-shaped relationship to well position. Bottom row: Data with strong systematic errors having a linear relationship to column number.

A widely used approach today is to design plate layouts manually in order to simplify for human interaction; e.g. placing controls in the outer-most wells (border layout) and distributing the samples following patterns that are easy to design and to pipette manually [18, 19]. Indeed many researchers still use border layouts as they help reduce human pipetting errors, allow for straight-forward visualisation of results by humans, for example in the form of heat maps [12], and can be easily designed using pen and paper [20]. Yet border layouts can only be used to effectively identify and adjust for only a few plate effects [13, 14], such as linear relationships to rows or columns that affect the whole plate (Figure 1).

For large-scale experiments with microplates having 384 or more wells, human pipetting becomes infeasible and robots for liquid handling are necessary. In recent years, pipetting robots have become common in biomedical labs and they allow for fully flexible arrangements of controls and samples on plates, making randomised layouts more accessible. However, pure randomisation can still produce ineffective layouts, for example large areas of the plate might end up not having any control samples, making it difficult or even impossible to detect and correct errors in those areas [16, 21, 22]. Further, replicates placed in adjacent wells are then likely to be affected by the same plate effects. Not only is it a problem that they will be similarly biased, but it has also been shown that clusters of similar samples, including similar doses of the same compound as well as technical replicates, can affect the results of adjacent wells [12]. Consequently, plate designs that distribute both controls and samples in a effective way are needed in order to reduce unwanted bias, as well as to aid to detect and correct plate effects. We refer to such designs as *effective* layouts. Figure 1 (top row) displays examples of microplates with two strong systematic plate effects (bowl-shaped and linear gradient) and examples of how controls can be located using border, random and effective layouts.

Several plate layout editors are available, such as Brunn [23], FlowJo [24], Labfolder [25], PlateDesigner [26], and PlateEditor [27]. While some are able to generate randomised layouts, none of them have capabilities to generate effective layouts. There is, of course, the possibility of generating several random layouts and then evaluate them in order to select the best one [28], but that does not guarantee that effective plate layouts have been selected, regardless of how many layouts are generated.

In this manuscript we introduce an artificial-intelligence based model for designing effective microplate layouts that can easily be adapted for different experimental settings, and evaluate it for dose-response and screening applications. In order to simplify its usage, we developed a suite of tools (PLAID), including a web-app for easily designing effective microplate layouts, together with Python notebooks for simulating different experimental designs and allow for planning and designing effective experiments.

### Effective microplate layouts

Below are listed properties that, in many cases, are relevant to construct effective plate layouts. The list is not meant to be exhaustive, and should be adapted for specific applications and experimental settings.

#### Distribution of control samples

In order to maximise the usefulness of positive and negative control samples during normalisation, controls should be distributed evenly among the wells of the microplate. For example, we could constrain the number of controls on each microplate to be equally distributed among each of its four quadrants, that is, the difference in the number of controls between any two quadrants would be at most 1. Moreover, controls could also be evenly distributed across rows and columns, which would be particularly useful to detect and mitigate plate effects linked to row or column number. Furthermore, controls of the same type should ideally not be placed on adjacent wells. Whenever feasible, we would also want controls of any kind not to be placed in adjacent wells.

#### Distribution of samples

It has been shown that a well with a strong effect can affect the measured intensity of its neighbour wells [11], and in particular, due to grouped similar samples [12]. With the goal of mitigating such *grouping effects* we can, for instance, enforce that the replicates of a sample are placed on different rows and columns. Similarly, for specific kinds of experiment, such as a dose-response experiment [29], we could enforce that for each compound, the difference in the number of individual doses between any two rows, any two columns, or any four quadrant is at most 1. Spreading samples with different doses this way makes the design resilient towards errors that affect an entire row or column (such as a pipetting error); enough of the other doses will remain for sufficient quality for e.g. regression.

#### Edge effects

Edge or border effects are discrepancies between the centre and the outer wells of a microplate primarily caused by evaporation during incubation, and can greatly affect the results obtained from an experiment [30]. A common method to mitigate edge effects is to avoid having samples in the outermost rows and columns, and instead fill them with medium or buffer [31].

#### Empty wells

If not all wells will be used, the locations of the empty wells could be constrained in a manner similar to that of control samples so they are distributed across the plate. This way, empty wells can help avoid clusters of samples and controls.

#### Multi-plate experiments

Across all plates, controls could also be balanced between plate halves or quadrants. Moreover, we could balance the controls per row or column across all plates, that is, the difference between the number of controls in any two rows or columns across all plates is at most 1. Given enough control samples, this can help ensure that potential plate effects linked to any row or column will be detected, especially when the errors have been introduced consistently in all plates, for example by a malfunctioning dispensing equipment. The same constraints could also be applied to sample replicates across plates.

### Effective layouts with constraint programming

Above we introduced desired properties of effective plate layouts as a set of constraints. One option to satisfy these constraints would be to randomly generate microplate layouts until one that fits the criteria is found. While this in itself constitutes a non-trivial task, finding such layout could take an unreasonably long time, and if no layout fulfilling the criteria exists, this program would never finish. A more efficient and natural solution is to frame our characterisation of effective microplate layouts as a constraint satisfaction problem (CSP): we view each well of each plate as a variable whose value represents its content and desirable properties of a layout as constraints. *Constraint programming* (CP) is a subarea of artificial intelligence that offers a flexible framework for solving constraint satisfaction problems that has seen large adoption in various fields (see Methods section). The general idea behind CP is that a CSP can be modelled as a conjunction of high-level constraints on variables ranging over initial domains, and then said model is given to a general-purpose constraint solver which performs a combination of intelligent reasoning and systematic search in order to find constraint-satisfying domain values for the variables. In this project we implement a constraint model that generates effective plate layouts for two different applications: dose-response and screening experiments.

## Effective layouts lead to more accurate results in dose-response experiments

Dose-response experiments attempt to evaluate the effect of a substance in a specific assay at increasing concentrations [29]. The effect can, in many cases, be estimated by fitting a sigmoid curve to the data points, and is frequently summarised by determining the half maximal inhibitory concentration (IC_50_), or the half maximal effective concentration (EC_50_). In order to evaluate the impact of different types of microplate layouts in dose-response experiments, we simulated a total of 43200 microplates for dose response experiments with border layouts, random layouts, and effective layouts generated using constraint programming and the constraints defined in Supplementary Listing 1. The experiments consisted of 20 compounds of varying potency in 6, 8, and 12 doses, and for 1, 2, and 3 replicates. Plate effects added had a relationship to column number or were bowl-shaped, both in medium and high strength. The data was normalised using linear regression in the case of border layouts, and LOESS regression for effective and random layouts, and four-parameter log-logistic (LL4) curves were fitted to the resulting data. Examples of the curves produced can be seen in Figure 2a and Supplementary Figure 1. For a complete description of the experiment, see Methods section.

**Figure 2:**
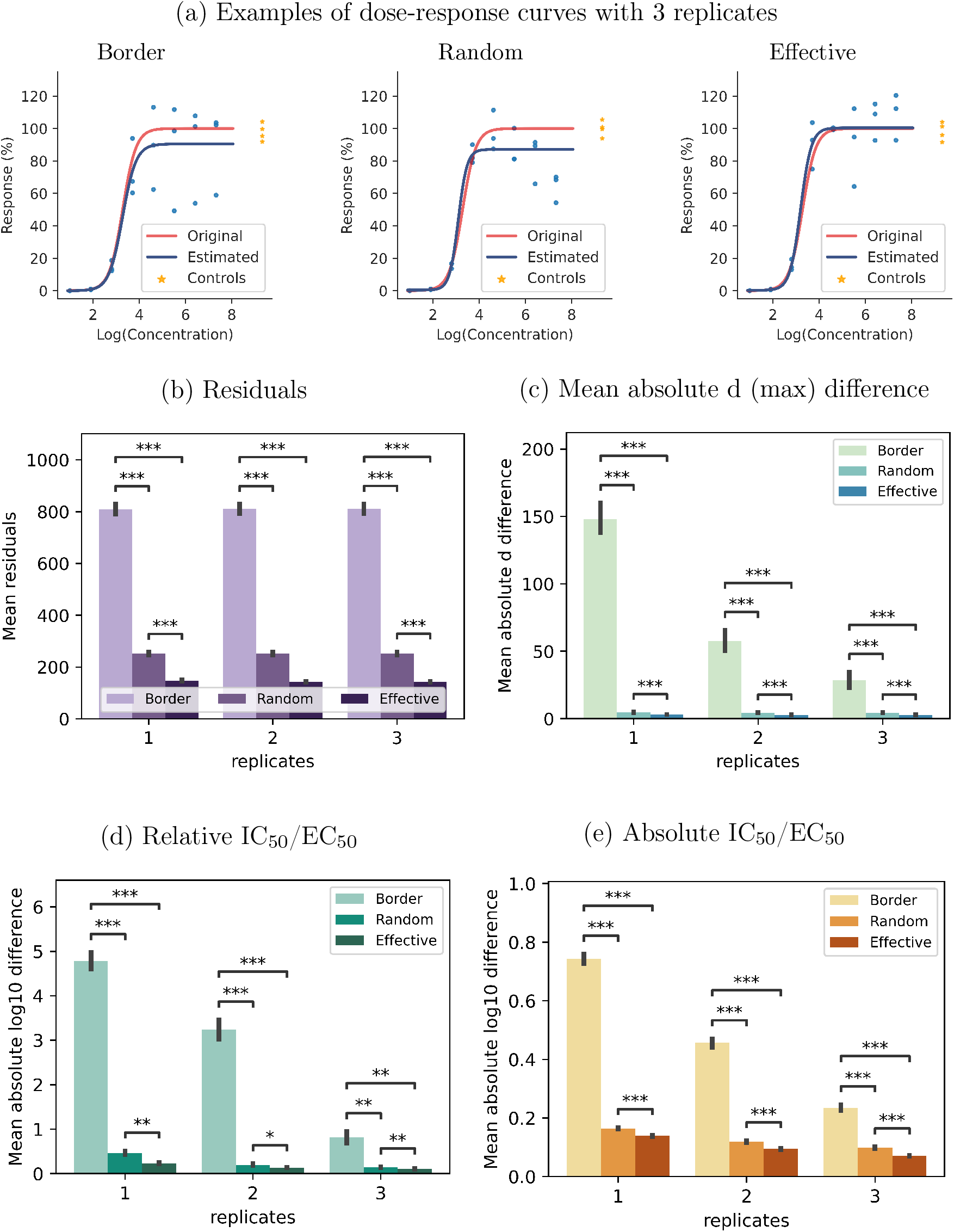
Comparison between expected and obtained values for dose response curves with 8 doses, 1, 2, or 3 replicates, 20 negative controls on 384-well plate, and strong plate effects with a linear relationship to column number on the right side of the plate. * indicates *p* < 10^−4^, ** indicates *p* < 10^−12^, *** indicates *p* < 10^−43^.

Figure 2b and Supplementary Figure 12 show the mean squared error (MSE) of the residuals calculated with respect to the dose-response curves used to generate the data. It is evident that, after error correction using LOESS regression and normalising to the mean of the negative controls, effective layouts lead to statistically significant smaller MSE than other types layouts (*p* < 10^−4^ for all pairwise comparisons, t-test). That is, the data obtained using our effective layouts is much closer to their expected values, than the data obtained when using either random and border layouts.

It is standard practice to discard dose-response curves that are considered to have low quality, for example, curves where more than 20% of the variability is unexplained by the curve fit, that is, with *R*^2^ < 0.8 [15]. In general, our effective layouts lead to a higher percentage of high-quality curves, as can be seen in Supplementary Figures 13–15. For example, in the case of experiments with 8 doses and 3 replicates, and strong plate effects with a linear relationship to column number on the right-half side of the plate, all curves generated using our effective layouts have an *R*^2^ ≥ 0.8, while only 94% of the curves generated using random layouts and 70% of the curves generated using border layouts have a good curve fit with *R*^2^ ≥ 0.8. Moreover, there is a significant difference between the various types of layouts when calculating the absolute difference between the maximum value of the expected and obtained curves as can be seen in Figure 2c and Supplementary Figures 2–4.

Estimated relative IC_50_/EC_50_ values from the data shows a significant difference between using an effective layout compared to using either a random or a border layout Figure 2d, regardless of the number of replicates used (*p* < 10^−4^ for all pairwise comparisons, t-test). In fact, we obtained a smaller MSE and a smaller standard deviation using 2 replicates and effective layouts than using 3 replicates and random layouts (Table 1). Similar results are obtained for other strengths of plate effects, as well as when using 6 or 12 doses (see Supplementary Figures 6 and 7).

**Table 1:** Mean log_10_ difference and standard deviation for the obtained relative IC_50_/EC_50_ and absolute IC_50_/EC_50_ for dose response curves with 8 doses, 1, 2, or 3 replicates, 20 negative controls on 384-well plate, and strong plate effects with a linear relationship to column number on the right side of the plate.

| Measurement | Layout | 1 replicate | 2 replicates | 3 replicates |
| --- | --- | --- | --- | --- |
| Relative IC<sub>50</sub>/EC<sub>50</sub> | Border | 4.78 ± (11.11) | 3.24 ± (9.17) | 0.81 ± (4.6) |
|  | Random | 0.47 ± (2.58) | 0.2 ± (0.86) | 0.14 ± (0.2) |
|  | Effective | 0.23 ± (0.94) | 0.13 ± (0.15) | 0.1 ± (0.13) |
| Absolute IC<sub>50</sub>/EC<sub>50</sub> | Border | 0.74 ± (1.01) | 0.46 ± (0.61) | 0.23 ± (0.35) |
|  | Random | 0.16 ± (0.15) | 0.12 ± (0.11) | 0.1 ± (0.09) |
|  | Effective | 0.14 ± (0.12) | 0.09 ± (0.08) | 0.07 ± (0.06) |

For estimated absolute IC_50_/EC_50_ values, Figure 2e shows that there is a significant difference between using an effective layout and either a random or a border layout regardless of the number of doses and replicates used (*p* < 10^−4^ for all pairwise comparisons, t-test). Similar results are obtained for other plate-effect strengths, and number of doses (see Supplementary Figures 8–10). Also note that it is not always possible to estimate the absolute IC_50_/EC_50_. For example, in the case of experiments with 8 doses and 1 replicate, the absolute IC_50_/EC_50_ of almost 1% of the curves could not be estimated when using border layouts in the presence of strong bowl-shaped effects. This number grows to 13.4% when the negative controls are not included as data points.

## Effective layouts improve sensitivity and reduce the risk of inflated quality assessment scores in screening experiments

Screening experiments attempt to identify hits from a large number of samples for further analysis [13, 14]. In order to evaluate the impact of different types of microplate layouts in screening experiments, we simulated 6480 384-well microplates using all combinations of: 40 layouts of each type of design, with either 8, 10, or 20 negative controls, 3 strength levels of bowl-shaped plate effects, and 6 hit percentages, namely 1%, 5%, 10%, 20%, 30%, and 40% of hits per plate. Each compound appears only once (1 replicate) and hits were randomly distributed on the plates. The results were adjusted using linear regression for border layouts, and LOESS regression for effective and random layouts. For a complete description of the experiment, see Methods section. Figures 3a-3c show examples of simulated screening data after error correction and normalisation in the presence of mild bowl-shaped plate effects. Figures 3f, 3g, and Supplementary Figures 22 and 23 show that, regardless of the number of negative controls used and hit rate, the use of effective layouts results in higher sensitivity (true positive rate) and yields statistically significant higher AUC (area under the curve) values with a smaller variance (Supplementary Tables 2 and 3; *p* < 10^−4^ for all pairwise comparisons, t-test).

**Figure 3:**
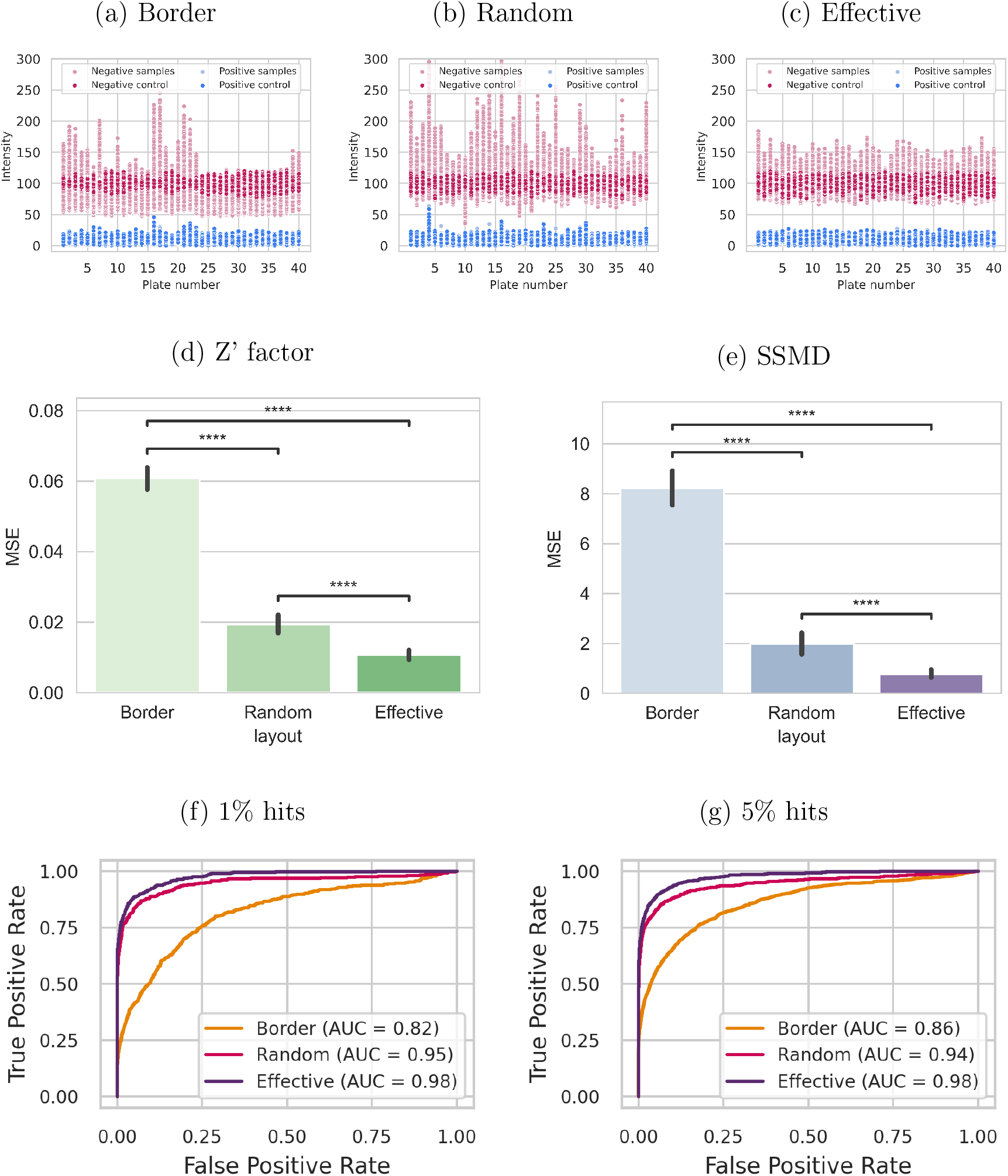
Screening experiments consisting of 40 microplates for each type of layout, each with 10 positive and 10 negative controls and only 1 replicate per compound. Hits are randomly distributed on each plate. (a)-(c): Simulated data after error correction and normalisation in the presence of mild bowl-shaped plate effects. MSE between expected and obtained (d) Z’ factor and (e) SSMD in the presence of mild bowl-shaped plate effects. The expected values were calculated using plates randomly filled with 50% positive controls and 50% negative controls, constituting the optimal values obtainable by these metrics. (f) and (g): ROC curves for experiments with varying hit rates in the presence of strong bowl-shaped plate effects.

Standard quality assessment metrics for microplate experiments include Z’ factor and SSMD [14], where low-quality-plate results are indicated by low metric scores, but where high scores are not a guarantee for high-quality results on the plate. The main reason for this is that both Z’ factor and SSMD only take into account positive and negatives controls regardless of their physical location on the plate, and with a sub-optimal layout these metrics might not accurately capture the real plate effects. In order to analyse the effect of different layouts on quality metrics, we calculated the expected values for both Z’ factor and SSMD using whole plates filled with 50% positive controls and 50% negative controls, constitut-ing the optimal quality values obtainable by these metrics. We then compared the resulting values against the same metrics being calculated using only a subset of the controls on the plate according to border, random, and effective layouts. Figures 3d and 3e show that for both the Z’ factor and SSMD, the estimates obtained using effective layouts yield a quality metric value that is closer to the expected value when compared to random and border layouts. This difference is always statistically significant as long as there is some degree of a plate effect (Supplementary Figures 24 and 25).

## The PLAID software suite

In order to make our method easily accessible, we developed PLAID (Plate Layouts using Artificial Intelligence Design), a suite of tools that can be used to design and evaluate microplate layouts under a wide range of conditions.

### The PLAID reference constraint model

We implemented a constraint model comprising the constraints described here using MiniZinc [32]. Advanced users can interact with and personalise the model by adding or removing constraints, which can be ran using the MiniZinc IDE, scripts, or command line. It is also possible to incorporate the model into existing workflows, for example, with the help of the MiniZinc Python package. Instructions on how to run our MiniZinc model using the command line or the MiniZinc IDE are available at https://github.com/pharmbio/plaid.

### The PLAID plate design tool

In order to ease the use of the PLAID constraint model, we developed an interactive web interface available at https://plaid.pharmb.io/ that allows for specifying experimental details and generating layouts (Supplementary Figure 26). The experimental design (e.g. selection of samples, concentrations, etc) can be downloaded from the web interface in a JSON format that can be later uploaded into the website in order to create more plate designs for the same experiment or as a base for new experiments. The produced layouts generated by the PLAID constraint model can be visualised within the web interface (Supplementary Figures 27 and 28), and downloaded in CSV, and JSON file formats, as well as an image (Supplementary Figure 29). Produced layouts in JSON format can be reuploaded into the website to use the visualisation features (Supplementary Figures 27 and 28). Examples of both experimental settings and layouts, as well as convenience methods for translating layouts into specific formats to be directly used in ECHO and I.DOT compound dispensing robots, are available on GitHub at https://github.com/pharmbio/plaid.

### The PLAID analysis and visualisation notebooks

Experiment designs can substantially vary, and no one-solution-fits-all exists. Different assays, laboratory conditions, equipment, etc, lead to different types and strengths of plate effects that affect experiments. We developed in Python a library of parametric plate effects, a library of plate normalisation and error correction functions, dose response and high throughput screening simulations, as well as visualisation functionality. This library can be used, for example from within Python notebooks, to evaluate different experimental designs, such as to explore the effect of varying the number of controls, doses, replicates, etc, before selecting the appropriate design.

## Discussion

Maximising the conclusions that can be drawn from data is the key objective when planning and carrying out biomedical experiments. With microplates becoming a standard platform to realise multiple-sample experiments, designing the physical layout of experiments and carrying out adequate data processing is essential to ensure high-quality data. Further, being able to minimise the number of control samples, replicates, or doses per sample can have a significant impact in terms of time, costs, and number of samples evaluated in any type of experiment. Research into data normalisation methods is an active subject, which has been especially important in omics research in the life science domain that, in many cases, can observe large variations between different labs, batches, and experimental settings. However, most normalisation techniques assume randomisation in the experimental design. We here show that randomising the physical locations of control samples can be sub-optimal, and that effective layouts generated using a constraint programming model are generally superior.

For dose-response experiments, effective layouts lead to significantly better approximations of curves when compared to random layouts, or especially the more traditional border layouts, on 384-well plates (Figure 2d). In fact, all curves in our experiments using effective layouts have an *R*^2^ *>* 0.8, implying that fewer approximation curves have to be discarded in experiments. Effective layouts also lead to significantly smaller MSE and standard deviation (Table 1 and Figure 2b,c) when estimating relative and absolute IC_50_/EC_50_ for dose-response curves. For screening experiments, the effect of plate layouts vary with the experiment, depending on the number of control samples, the expected hit rate, and the strength of systematic errors. Our experiments demonstrate that for experiments with strong bowl-shaped errors, effective layouts have a significantly higher sensitivity compared with random and border layouts with as low as 1% hit rates – even when samples do not have any replicates (Figure 3f). With lower systematic errors, the impact of using effective layouts is smaller but still relevant, especially for experiments having higher hit rates as shown by Mpindi et al. [13] (Supplementary Figures 22 and 23). These results underline the value of experimental design and physical placement of samples also for screening experiments. Effective layouts also reduce the risk of obtaining an inflated score from plate quality assessment metrics such as Z’ factor and SSMD, as the values for such metrics when calculated on effective layouts are on average closer to the expected (optimal) metric value compared to random and border layouts.

Simulating multiple scenarios allows for evaluating and comparing different experimental parameters, such as the effect of the number of replicates versus the number of concentrations per sample. This can, for example, be carried out and visualised using the provided PLAID notebooks. Our results show that in common dose-response experiments, effective layouts can lead to a reduction in the number of replicates while maintaining a higher confidence in the estimated IC_50_/EC_50_ (Figure 2b,c). We also observe, inline with the recommendations in [33], that replicates do improve precision, but not enough to address systematic bias. In general, adding more doses had a higher impact in the estimations than adding more replicates, regardless of the layout. In particular, effective layouts generally lead to more accurate results even with fewer replicates or fewer doses. For example, we obtained more accurate estimations of absolute IC_50_/EC_50_ for experiments with 8 doses and 2 replicates using our effective layouts than with 8 doses and 3 replicates using random layouts (see Figure 2e and Supplementary Figures 8– 10). Moreover, we also obtained more accurate estimations of absolute EC_50_/IC_50_ for experiments with 8 doses and 3 replicates using our effective layouts, compared to 12 doses and 3 replicates using random layouts (see Supplementary Figures 8– 10).

The benefits and limitations of effective microplate layouts are tightly coupled to the use and impact of methods for data normalisation, that in turn are dependent on the use of, and sufficient number of, control samples. Further, a multi-plate experiment also offers more opportunities for finding effective layouts. In our work we focused on LOESS normalisation [34], which is a widely used normalisation technique that is robust towards different types of plate and experimental effects. The error model used in this work is based on the one proposed by Zhang et al. [11], but the intensity and type of plate effects observed might differ depending on factors such as the type of experiment, laboratory facilities, and temperature, among others. A key advantage of using a constraint model for designing layouts is that such parameters can be easily adjusted, due to the declarative nature of the model, and evaluated using the provided PLAID Python notebooks. We have put together a suite of constraints that are widely useful, but the final selection of constraints is up to the scientists planning experiments and it is easy to e.g. remove constraints such as ‘no samples in outer wells’ if they are not desirable. From a practical perspective, such as when pipetting manually, it can be beneficial to use the same sample only on one plate. For automated liquid handling instruments, the number of plates (and hence source samples) accessible can have an impact. These scenarios are not covered in this study, but there is no hinder to also implement such constraints into a plate layout model.

### Iterative experimentation

AI methods such as unsupervised and supervised learning are nowadays widely used to analyse the results from large-scale experiments using microplates. There are also emerging approaches to sequentially plan a series of experiments to systematically improve the accuracy of AI models [35, 36]. Data-centric AI is a concept that proposes to shift from the current practice of having a set of fixed data and then spending much time and effort to fine-tune a machine learning model, to instead focus on an iterative approach to optimise the data used to train the model [37]. This methodology fundamentally builds on the proposition that high-quality data is better than just more data, something that for a long time has been argued in traditional experimental design guidelines [33]. Selecting the next batch of experiments is however non-trivial, and autonomous decision-making is currently an active research field with autonomous vehicles as a big driver. Active machine learning is one approach to select new experiments which, combined with robotics, has the potential to automate scientific discoveries [38–41].

Figure 4 shows how the PLAID suite supports iterative experimentation. The first step constitutes an initial decision on samples, replicates, controls, etc., and its definition in a declarative file format for microplate experiments (Supplementary Section 1). This experiment-definition file is then input to the PLAID plate design tool that applies the constraint model to generate effective plate layouts. The produced layouts can then be evaluated towards simulated experiments defined by different error parameters, which over time can be tailored to particular experimental and laboratory setups. Based on the outcome of the simulations, the experiment design might be revised and new plate layouts can be generated. When a decision is made to accept the layouts, these can be translated to custom formats that can be read by lab instruments. We provide translations for two common chemical dispensing instruments (ECHO and I.DOT), but it is straight-forward to create more adaptors for other instruments. Accepting plate layouts can be done manually by humans, or autonomously using an algorithm. If the data acquisition and analysis from the physical experiments can be automated, then only the decision making on the next round of experiments remains. Given that autonomous decision making has been implemented in order to select the next experiment, it can be defined in the PLAID file format for microplate experiments, closing the loop for the next experiment iteration. We speculate that such automated and iterative scientific experiments will be increasingly common in the future, and that PLAID, given its flexibility due to the declarative nature of constraint modelling, its open source implementation, and associated tools for easy integration and visualisation, is a compelling model and architecture.

**Figure 4:**
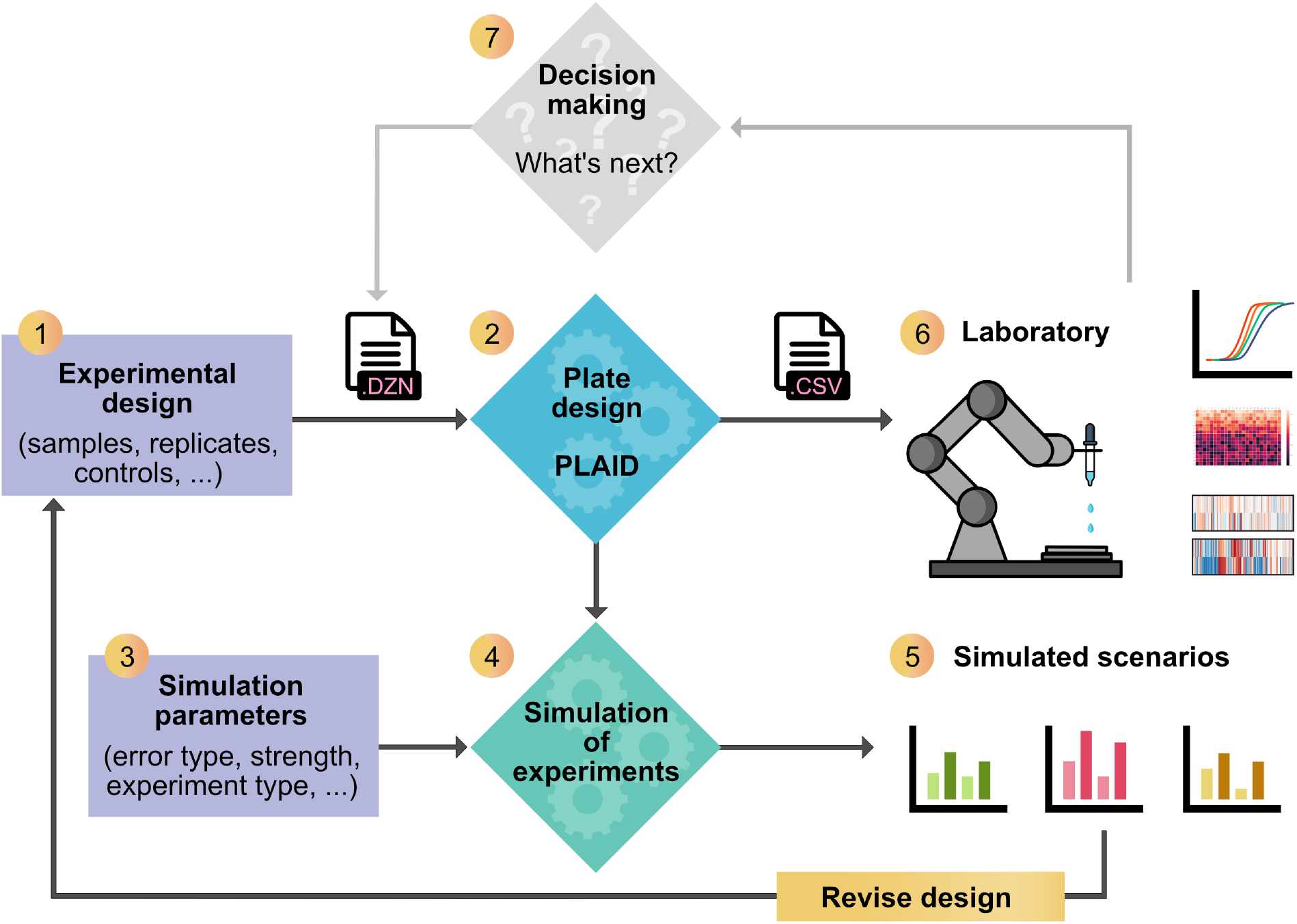
Overview of the PLAID ecosystem. Experiment design (1) comprises the selection of samples, replicates, concentrations, etc, and can be defined in a configuration file or via the PLAID Plate Design web interface, and then the PLAID constraint model is called to run the optimisation and generate the resulting layouts (2). In order to evaluate the design, simulation parameters such as errors need to be defined (3) and then a simulated experiment using the layouts can be carried out in the PLAID Analysis and Visualisation Notebooks (4). Different scenarios can be compared, such as different numbers of concentrations and replicates, and designs may then be revised accordingly. When an acceptable design has been generated, it can be used to drive liquid handling instruments, such as automated pipette robots (6). After the experiment is performed and analysed, a decision can be made on subsequent experiments e.g. confirm findings, re-run failed samples, evaluate more concentrations, etc. (7). Implementing automated decision making that defines the next experiment in the PLAID configuration file format enables autonomous sequential experimentation.

## Conclusions

We identified properties of effective microplate layouts and used artificial intelligence in the form of constraint programming to build a model that is capable of generating such layouts, and evaluated their effect on normalisation in common experiment settings involving multi-plates. We demonstrate that effective layouts are superior to random layouts for illustrative dose-response and screening experiments, generating more robust results and data with lower variance and higher sensitivity. The software suite PLAID makes the method easily available, allows for decision aid in experiment design to select e.g. number of doses and replicates, and is prepared for integration into closed-loop systems. Examples of studies where PLAID has been used include [42–44].

## Methods

### Constraint programming

Constraint programming (CP) [45] is a form of artificial intelligence used for modelling and solving combinatorial problems, which is currently successfully used in many real-world application areas such as scheduling [46–48], decision support [49], and packing [50]. Solving a *combinatorial problem* involves finding an assignment for a discrete, finite set of objects (decision variables) that satisfies a given set of conditions (constraints). The general idea behind constraint programming is that the user specifies the constraints that should hold among decision variables and a general-purpose constraint solver is used to find a solution. That is, the user specifies the problem without having to specify how to find a solution. For example, consider our microplate layout design problem. Each unknown in the problem, namely the content of each well on each plate, is a decision variable. Each decision variable *V*_*i*_ can take values in a given domain, denoted dom(*V*_*i*_). In our microplate layout design problem, the domain of each decision variable is the set of possible substances to place on a well, i.e. a given compound at a certain concentration, a positive control, etc. Moreover, problem solutions are distinguished from non-solutions by constraints, which are the limitations to the values that the decision variables can take simultaneously. In this context, a constraint is, for example, a limitation that controls of the same type cannot be placed in contiguous wells.

In order to find a solution for a given problem, a constraint solver first removes infeasible values from the domains of the variables by applying inference methods, which is known in the literature as *propagation*. Then, the search for a feasible solution is performed in a branch-and-bound fashion: the left-most branch corresponds to a sub-problem that is created by assigning a value *v* ∈ dom(*V*_*i*_) to a variable *V*_*i*_. If the sub-problem turns out to be infeasible, a backtracking mechanism is used to try other sub-problems where the additional constraint dom(*V*_*i*_) ≠ *v* is added.

In general, constraint satisfaction problems are specified by data-independent models written in a modelling language such as AMPL [51], Essence [52], MiniZinc [32], or OPL [53].

### Constraint model implementation

We implemented a constraint model representing the microplate layout design problem in MiniZinc [32] and used Gecode [54] as the back-end constraint solver. One of the many advantages of using MiniZinc is that only very minor modifications, if any, would be needed to use another constraint solver. Examples of constraints included in our model together with their representation in MiniZinc can be seen in Figure 5. For a full list of all constraints defined in this study, see Supplementary Listing 1.

**Figure 5:**
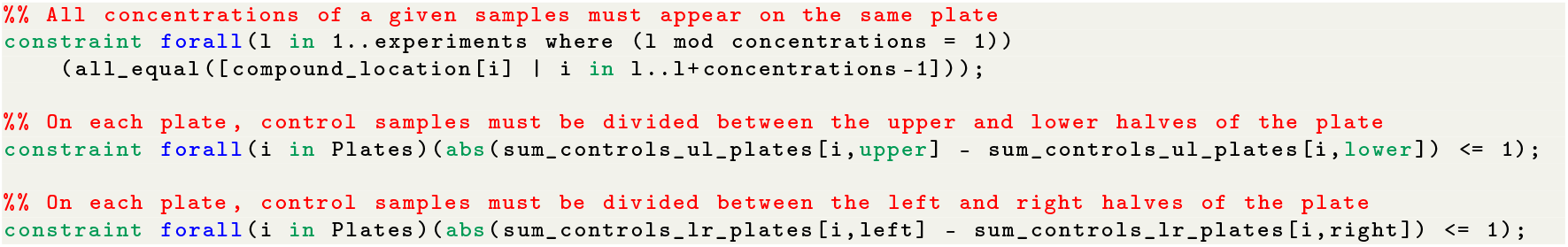
Examples of constraints in MiniZinc syntax

On top of including all the desirable properties of effective microplate layouts, we have chosen to include other constraints that are needed for practical matters. For example, we enforce that for each sample, all concentration levels of a given replica must appear on the same plate. Technical replicates of a sample can be chosen to appear on the same plate, on a different plate, or a mixture of both. We have also included the dimensions of the microplate as parameters in terms of number of rows and columns, allowing the use of any kind of plate size. Finally, it is also possible to specify how many rows and columns should be left empty on the border of every microplate in order to mitigate edge effects.

### Dose-response experiments

We simulated multiple scenarios for dose-response experiments according to [15]. The following scenarios were considered: all combinations of compounds having: (i) a sigmoid curve with slopes of 0.5, 1, 1.5, and 2; (ii) 6 concentrations with a dilution factor of 18, 8 concentrations with a dilution factor of 8, and 16 concentrations with a dilution factor of 4; and (iii) 1, 2, and 3 replicates per compound. Without loss of generality, for every compound the bottom of the curve was set to 0%, and the top of the curve was set to 100%. Fixing the top and bottom of the curve at these values makes the assumption that if a sufficient number of concentrations were to be used, a complete dose–response curve would be generated. To generate the sigmoid curves corresponding to each compound the only parameter remaining to be specified is the EC_50_/IC_50_. We generated curves with EC_50_/IC_50_ values ranging from 1 to 96 to simulate compounds having all kinds of potency. The highest concentration was arbitrarily set to 100 µM. For each test concentration, the replicates were generated by adding a random value within ±1% to the value sampled from the curve in order to represent a very small error in measurement between wells having the same compound in the same concentration.

Border layouts were designed by placing 20 negative controls in columns 2 and 23, and all other samples were placed horizontally from top to bottom. Random layouts were generated using the Python random package. Effective microplate layouts were generated using our constraint programming model implemented in MiniZinc [32]. The Python functions and MiniZinc model used to generate the plates and the resulting layouts are available on https://github.com/pharmbio/plaid. We then applied the same plate effect to every plate having either: (i) a bowl-shape relationship to well position, or (ii) a linear relationship to column number on the right-hand side of the plate. Strong plate effects are designed according to the examples in [11, 14], while medium plate effects are half way between no-effect and a strong plate effect. After applying plate effects, we adjust the data using linear regression in the case of border layouts, and LOESS regression as implemented in [55] for the rest, and normalised the data as a percentage of the mean of the negative controls. Finally, we estimate the relative and absolute EC_50_/IC_50_ using the `curve_fit` function of the `scipy` Python library, which uses the Trust Region Reflective algorithm. For each dose-response curve, we calculated the absolute value of the difference between the log_10_ of the true and the estimated EC_50_/IC_50_ values. Moreover, for every measurement, we calculated the difference with respect to both the expected (true) value as well as with respect to the estimated curves.

### Screening experiments

We simulated screening experiments with 40 384-well microplates, each of which contained either (i) 8 positive controls and 8 negative controls, (ii) 10 positive controls and 10 negative controls, or (iii) 10 positive controls and either 20 negative controls. The remaining wells contained random samples with hit-rates of 1%, 5%, 10%, 20%, 33%, and 40%. We then applied various strengths of bowl-shaped effects to every plate. Strong plate effects are designed according to the examples in [11, 14], while medium plate effects are half way between no-effect and a strong plate effect. After applying the plate effect, we calculated the raw Z’ factor and the raw SSMD of each plate. For normalisation we used linear regression in the case of border layouts, and loess regression (as implemented in [55]) in the case of random and effective layouts, and scaled the data as a percentage of the average of the nearest negative controls. Both linear and loess regression are performed based on negative controls only, without assuming a low hit-rate. We used the error-corrected data to calculate the final Z’ factor and the SSMD of each microplate. Finally, we used the `sklearn` Python library to calculate the resulting ROC curves and AUC values.

## Data availability

The Python libraries and notebooks developed for the analysis, the experimental results, together with the specific microplate layouts tested are available at https://github.com/pharmbio/plaid.

## Code availability

All source code for PLAID, including our constraint model, libraries and Python notebooks for simulating, evaluating and visualising experiments, as well as scripts for layout translations are open source and publicly available at https://github.com/pharmbio/plaid. A managed service for public consumption of the web interface is available at https://plaid.pharmb.io/, and its Docker image is available at https://github.com/pharmbio/plaid-gui.

## Acknowledgements

This project received funding from the Swedish Research Council (grants 2020-03731 and 2020-01865), FORMAS (grant 2018-00924), the Swedish Foundation for Strategic Research (grant BD15-0008SB16-0046), and the Swedish strategic research programme eSSENCE.

We thank Wesley Schaal for constructive feedback on manuscript, Markus Lucero and Travis Persson for contributions to the PLAID web interface, Polina Georgiev for constructive feedback on microplate layouts, Ebba Bergman for constructive feedback on figures, and Gustav Björdal for constructive feedback on the constraint model.

## Author information

### Contributions

MAFR and OS: Project conceptualisation; MAFR: Method development and implementation, tool development; MAFR, OS, JCP: Analysis and interpretations, manuscript preparation.

## References

[1] Jessica Vamathevan, Dominic Clark, Paul Czodrowski, Ian Dunham, Edgardo Ferran, George Lee, Bin Li, Anant Madabhushi, Parantu Shah, Michaela Spitzer, and Shanrong Zhao. Applications of machine learning in drug discovery and development. Nature Reviews Drug Discovery, 18(6):463–477, June 2019.

[2] Ahmed M. Alaa, Deepti Gurdasani, Adrian L. Harris, Jem Rashbass, and Mihaela van der Schaar. Machine learning to guide the use of adjuvant therapies for breast cancer. Nature Machine Intelligence, 3(8):716–726, August 2021.

[3] Brodie Fischbacher, Sarita Hedaya, Brigham J. Hartley, Zhongwei Wang, Gregory Lallos, Dillion Hutson, Matthew Zimmer, Jacob Brammer, and Daniel Paull. Modular deep learning enables automated identification of monoclonal cell lines. Nature Machine Intelligence, 3(7):632–640, July 2021.

[4] James Lu, Brendan Bender, Jin Y. Jin, and Yuanfang Guan. Deep learning prediction of patient response time course from early data via neural-pharmacokinetic/pharmacodynamic modelling. Nature Machine Intelligence, 3(8):696–704, August 2021.

[5] Alex Mattenet, Ian Davidson, Siegfried Nijssen, and Pierre Schaus. Generic Constraint-based Block Modeling using Constraint Programming. Journal of Artificial Intelligence Research, 70:597–630, February 2021.

[6] Luisa Franchina and Federico Sergiani. High Quality Dataset for Machine Learning in the Business Intelligence Domain. In Yaxin Bi, Rahul Bhatia, and Supriya Kapoor, editors, Intelligent Systems and Applications, Advances in Intelligent Systems and Computing, pages 391–401, Cham, 2020. Springer International Publishing.

[7] Yuzhen Lu and Sierra Young. A survey of public datasets for computer vision tasks in precision agriculture. Computers and Electronics in Agriculture, 178:105760, November 2020.

[8] Christopher J. Williams, David C. Richardson, and Jane S. Richardson. The importance of residue-level filtering and the Top2018 best-parts dataset of high-quality protein residues. Protein Science, n/a(n/a), 2021.

[9] Michal Alexovič, Pawel L. Urban, Hadi Tabani, and Ján Sabo. Recent advances in robotic protein sample preparation for clinical analysis and other biomedical applications. Clinica Chimica Acta, 507:104–116, August 2020.

[10] Ji-Hu Zhang, Thomas D. Y. Chung, and Kevin R. Oldenburg. A Simple Statistical Parameter for Use in Evaluation and Validation of High Throughput Screening Assays. Journal of Biomolecular Screening, 4(2):67–73, April 1999.

[11] Xiaohua Douglas Zhang. Experimental Designs. In Optimal High-Throughput Screening: Practical Experimental Design and Data Analysis for Genome-Scale RNAi Research, pages 13–26. Cambridge University Press, Cambridge, 2011.

[12] Amanda Birmingham, Laura M. Selfors, Thorsten Forster, David Wrobel, Caleb J. Kennedy, Emma Shanks, Javier Santoyo-Lopez, Dara J. Dunican, Aideen Long, Dermot Kelleher, Queta Smith, Roderick L. Beijersbergen, Peter Ghazal, and Caroline E. Shamu. Statistical methods for analysis of highthroughput RNA interference screens. Nature Methods, 6(8):569–575, August 2009.

[13] John-Patrick Mpindi, Potdar Swapnil, Bychkov Dmitrii, Saarela Jani, Khalid Saeed, Krister Wennerberg, Tero Aittokallio, Päivi Östling, and Olli Kallioniemi. Impact of normalization methods on high-throughput screening data with high hit rates and drug testing with dose–response data. Bioinformatics, 31(23):3815–3821, December 2015.

[14] Xiaohua Douglas Zhang. Novel Analytic Criteria and Effective Plate Designs for Quality Control in Genome-Scale RNAi Screens. Journal of Biomolecular Screening, 13(5):363–377, June 2008.

[15] J. L. Sebaugh. Guidelines for accurate EC50/IC50 estimation. Pharmaceutical Statistics, 10(2):128–134, 2011.

[16] Christopher Roselle, Thorsten Verch, and Mary Shank-Retzlaff. Mitigation of microtiter plate positioning effects using a block randomization scheme. Analytical and Bioanalytical Chemistry, 408(15):3969–3979, June 2016.

[17] Juan C. Caicedo, Sam Cooper, Florian Heigwer, Scott Warchal, Peng Qiu, Csaba Molnar, Aliaksei S. Vasilevich, Joseph D. Barry, Harmanjit Singh Bansal, Oren Kraus, Mathias Wawer, Lassi Paavolainen, Markus D. Herrmann, Mohammad Rohban, Jane Hung, Holger Hennig, John Concannon, Ian Smith, Paul A. Clemons, Shantanu Singh, Paul Rees, Peter Horvath, Roger G. Linington, and Anne E. Carpenter. Data-analysis strategies for image-based cell profiling. Nature Methods, 14(9):849–863, September 2017.

[18] Stacey L. Brower, Jeffrey E. Fensterer, and Jason E. Bush. The ChemoFx ®Assay: An Ex Vivo Chemosensitivity and Resistance Assay for Predicting Patient Response to Cancer Chemotherapy. In Gil Mor and Ayesha B. Alvero, editors, Apoptosis and Cancer: Methods and Protocols, Methods in Molecular Biology™, pages 57–78. Humana Press, Totowa, NJ, 2008.

[19] Steven P. Williams, Cathryn M. Gould, Cameron J. Nowell, Tara Karnezis, Marc G. Achen, Kaylene J. Simpson, and Steven A. Stacker. Systematic highcontent genome-wide RNAi screens of endothelial cell migration and morphology. Scientific Data, 4(1):170009, March 2017.

[20] Brian Connelly. Plotting Microtiter Plate Maps. https://bconnelly.net/posts/plotting-microtiter-plate-maps/, May 2014.

[21] Hélène Borges, Anne-Marie Hesse, Alexandra Kraut, Yohann Couté, Virginie Brun, and Thomas Burger. Well Plate Maker: A user-friendly randomized block design application to limit batch effects in largescale biomedical studies. Bioinformatics (Oxford, England), page btab065, February 2021.

[22] Bram Burger, Marc Vaudel, and Harald Barsnes. Importance of Block Randomization When Designing Proteomics Experiments. Journal of Proteome Research, 20(1):122–128, January 2021.

[23] Jonathan Alvarsson, Claes Andersson, Ola Spjuth, Rolf Larsson, and Jarl ES Wikberg. Brunn: An open source laboratory information system for microplates with a graphical plate layout design process. BMC Bioinformatics, 12(1):179, May 2011.

[24] FlowJo, LLC. https://www.flowjo.com/.

[25] Well Plate Templates. https://www.labfolder.com/well-plate-templates/.

[26] Maria Suprun and Mayte Suárez-Fariñas. PlateDesigner: A web-based application for the design of microplate experiments. Bioinformatics (Oxford, England), 35(9):1605–1607, May 2019.

[27] Vincent Delorme, Minjeong Woo, Virginia Carla de Almeida Falcão, and Connor Wood. PlateEditor: A web-based application for the management of multi-well plate layouts and associated data. PLOS ONE, 16(5):e0252488, May 2021.

[28] Lauren Schiff, Bianca Migliori, Ye Chen, Deidre Carter, Caitlyn Bonilla, Jenna Hall, Minjie Fan, Edmund Tam, Sara Ahadi, Brodie Fischbacher, Anton Geraschenko, Christopher J. Hunter, Subhashini Venugopalan, Sean DesMarteau, Arunachalam Narayanaswamy, Selwyn Jacob, Zan Armstrong, Peter Ferrarotto, Brian Williams, Geoff Buckley-Herd, Jon Hazard, Jordan Goldberg, Marc Coram, Reid Otto, Edward A. Baltz, Laura Andres-Martin, Orion Pritchard, Alyssa Duren-Lubanski, Kathryn Reggio, NYSCF Global Stem Cell Array Team, Lauren Bauer, Raeka S. Aiyar, Elizabeth Schwarzbach, Daniel Paull, Scott A. Noggle, Frederick J. Monsma, Marc Berndl, Samuel J. Yang, and Bjarki Johannesson. Deep learning and automated Cell Painting reveal Parkinson’s disease-specific signatures in primary patient fibroblasts. bioRxiv, page 2020.11.13.380576, November 2020.

[29] Laura N. Vandenberg. Chapter 7 - Low Dose Effects and Nonmonotonic Dose Responses for Endocrine Disruptors. In Philippa D. Darbre, editor, Endocrine Disruption and Human Health (Second Edition), pages 141–163. Academic Press, January 2022.

[30] Jean-Philippe Carralot, Arnaud Ogier, Annette Boese, Auguste Genovesio, Priscille Brodin, Peter Sommer, and Thierry Dorval. A novel specific edge effect correction method for RNA interference screenings. Bioinformatics, 28(2):261–268, January 2012.

[31] Betina Kerstin Lundholt, Kurt M. Scudder, and Len Pagliaro. A Simple Technique for Reducing Edge Effect in Cell-Based Assays. Journal of Biomolecular Screening, 8(5):566–570, October 2003.

[32] Nicholas Nethercote, Peter J. Stuckey, Ralph Becket, Sebastian Brand, Gregory J. Duck, and Guido Tack. MiniZinc: Towards a Standard CP Modelling Language. In Christian Bessière, editor, Principles and Practice of Constraint Programming –CP 2007, Lecture Notes in Computer Science, pages 529–543, Berlin, Heidelberg, 2007. Springer.

[33] U.S. Pharmacopeial Convention. USP_1032.pdf.

[34] WS Cleveland, E Grosse, and WM Shyu. Local regression models. chapter 8 in statistical models in s (jm chambers and tj hastie eds.), 608 p. Wadsworth & Brooks/Cole, Pacific Grove, CA, 1992.

[35] Natalie S Eyke, William H Green, and Klavs F Jensen. Iterative experimental design based on active machine learning reduces the experimental burden associated with reaction screening. Reaction Chemistry & Engineering, 5(10):1963–1972, 2020.

[36] Sean Ekins, Ana C Puhl, Kimberley M Zorn, Thomas R Lane, Daniel P Russo, Jennifer J Klein, Anthony J Hickey, and Alex M Clark. Exploiting machine learning for end-to-end drug discovery and development. Nature materials, 18(5):435–441, 2019.

[37] Lester James Miranda. Towards data-centric machine learning: a short review. ljvmiranda921.github.io, 2021.

[38] Gisbert Schneider. Automating drug discovery. Nature Reviews Drug Discovery, 17(2):97–113, 2018.

[39] Ross D King, Jem Rowland, Stephen G Oliver, Michael Young, Wayne Aubrey, Emma Byrne, Maria Liakata, Magdalena Markham, Pinar Pir, Larisa N Soldatova, Andrew Sparkes, Kenneth E Whelan, and Amanda Clare. The automation of science. Science, 324(5923):85–89, April 2009.

[40] Semion K Saikin, Christoph Kreisbeck, Dennis Sheberla, Jill S Becker, and Alán Aspuru-Guzik. Closed-loop discovery platform integration is needed for artificial intelligence to make an impact in drug discovery. Expert opinion on drug discovery, 14(1):1–4, 2019.

[41] Daniel Reker. Practical considerations for active machine learning in drug discovery. Drug Discovery Today: Technologies, 32:73–79, 2019.

[42] Ankit Gupta, Philip J. Harrison, Håkan Wieslander, Jonne Rietdijk, Jordi Carreras Puigvert, Polina Georgiev, Carolina Wählby, Ola Spjuth, and Ida-Maria Sintorn. Is brightfield all you need for mechanism of action prediction?, October 2022.

[43] Jonne Rietdijk, Tanya Aggarwal, Polina Georgieva, Maris Lapins, Jordi Carreras-Puigvert, and Ola Spjuth. Morphological profiling of environmental chemicals enables efficient and untargeted exploration of combination effects. Science of The Total Environment, 832:155058, August 2022.

[44] Guangyan Tian, Philip J. Harrison, Akshai P. Sreenivasan, Jordi Carreras Puigvert, and Ola Spjuth. Combining molecular and cell painting image data for mechanism of action prediction, October 2022.

[45] Stuart Russell and Peter Norvig. Artificial Intelligence: A Modern Approach. Pearson, 4th global edition edition, 2021.

[46] Ekaterina Arafailova, Nicolas Beldiceanu, Rémi Douence, Pierre Flener, María Andreína Francisco Rodríguez, Justin Pearson, and Helmut Simonis. Time-Series Constraints: Improvements and Application in CP and MIP Contexts. In Claude-Guy Quimper, editor, Integration of AI and OR Techniques in Constraint Programming, Lecture Notes in Computer Science, pages 18–34, Cham, 2016. Springer International Publishing.

[47] Sara Frimodig and Christian Schulte. Models for Radiation Therapy Patient Scheduling. In Thomas Schiex and Simon de Givry, editors, Principles and Practice of Constraint Programming, Lecture Notes in Computer Science, pages 421–437, Cham, 2019. Springer International Publishing.

[48] Tobias Geibinger, Florian Mischek, and Nysret Musliu. Investigating Constraint Programming for Real World Industrial Test Laboratory Scheduling. In Louis-Martin Rousseau and Kostas Stergiou, editors, Integration of Constraint Programming, Artificial Intelligence, and Operations Research, Lecture Notes in Computer Science, pages 304–319, Cham, 2019. Springer International Publishing.

[49] Saumya Bhatnagar, Akshat Kumar, and Hoong Chuin Lau. Decision making for improving maritime traffic safety using constraint programming. Proceedings of the Twenty-Eighth International Joint Conference on Artificial Intelligence 2019:Macau, August 10-16, pages 5794–5800, August 2019.

[50] Aline A. S. Leao, Franklina M. B. Toledo, José Fernando Oliveira, Maria Antónia Carravilla, and Ramón Alvarez-Valdés. Irregular packing problems: A review of mathematical models. European Journal of Operational Research, 282(3):803–822, May 2020.

[51] Robert Fourer, David M Gay, and Brian W Kernighan. A modeling language for mathematical programming. Management Science, 36(5):519–554, 1990.

[52] Alan M Frisch, Warwick Harvey, Chris Jefferson, Bernadette Martínez-Hernández, and Ian Miguel. Essence: A constraint language for specifying combinatorial problems. Constraints, 13(3):268–306, 2008.

[53] Pascal Van Hentenryck, Laurent Michel, Laurent Perron, and J-C Régin. Constraint programming in opl. In International Conference on Principles and Practice of Declarative Programming, pages 98–116. Springer, 1999.

[54] Gecode Team. Gecode: Generic constraint development environment, 2019. Available from http://www.gecode.org.

[55] Michele Cappellari, Richard M. McDermid, Katherine Alatalo, Leo Blitz, Maxime Bois, Frédéric Bournaud, M. Bureau, Alison F. Crocker, Roger L. Davies, Timothy A. Davis, P. T. de Zeeuw, Pierre-Alain Duc, Eric Emsellem, Sadegh Khochfar, Davor Krajnović, Harald Kuntschner, Raffaella Morganti, Thorsten Naab, Tom Oosterloo, Marc Sarzi, Nicholas Scott, Paolo Serra, Anne-Marie Weijmans, and Lisa M. Young. The ATLAS3D project – XX. Mass–size and mass–σ distributions of early-type galaxies: Bulge fraction drives kinematics, mass-to-light ratio, molecular gas fraction and stellar initial mass function. Monthly Notices of the Royal Astronomical Society, 432(3):1862–1893, July 2013.

