## Supplemental Material for "Designing microplate layouts using artificial intelligence"

### Designing microplate layouts using artificial intelligence: supplementary information

#### Listing 1: List of constraints

An implementation of these constraints using the MiniZinc constraint programming language can be found at `https://github.com/pharmbio/plaid`.

##### Listing 1.1: Control samples (both positive and negative)

- On each microplate:
  - the difference in the total number of control samples between the upper and lower halves of the microplate is at most 1.
  - the difference in the total number of control samples between the left and right halves of the microplate is at most 1.
  - the difference in the total number of control samples between any two quadrants is at most 1.
  - the difference in the total number of control samples between any two rows is at most 1.
  - the difference in the total number of control samples between any two columns is at most 1.
  - the difference in the number of control samples of each type between the upper and lower halves of the microplate is at most 1.
  - the difference in the number of control samples of each type between the left and right halves of the microplate is at most 1.
  - the difference in the number of control samples of each type between any two quadrants is at most 1.
  - the difference in the number of control samples of each type between any two rows is at most 1.

  - the difference in the number of control samples of each type between any two columns is at most 1.
  - control samples are not side by side, that is, a control sample does not have another control sample on the wells that are directly to its left or right, or above or below.
  - if the microplates are large enough (when the number of available wells in the microplate is at least 4 times larger than the number of control samples) then control samples of any kind are not in adjacent wells, that is, control samples of the same type are not placed left, right, above, bellow or diagonally to each other.
  - if the microplates are large enough (when the number of available wells in the microplate is at least 4 times the number of a given kind of control samples) then control samples of a each particular kind are not on adjacent wells, that is, control samples of the same type are not placed left, right, above, bellow or diagonally to each other.
  - if the microplates are large enough (when the number of available wells in the microplate is at least 9 times the number of a given kind of control samples) then control samples of a each particular kind have at least 2 wells in between.

- Across all microplates:
  - the difference in the total number of control samples between any two rows is at most 1.
  - the difference in the total number of control samples between any two columns is at most 1.

#### Listing 1.2: Samples

- On each microplate:
  - replicates of a sample are placed on different rows
  - replicates of a sample are placed on different columns
  - the difference in the total number of samples between the upper half and the lower half of the microplate is at most 1.
  - the difference in the total number of samples between the left and the right halves of the microplate is at most 1.
  - the difference in the total number of samples between any two rows is at most 1.
  - the difference in the total number of samples between any two columns is at most 1.
  - the difference in the total number of samples between the left half and the right half of every row is at most 1.
  - the difference in the total number of samples between the upper half and the lower half of every column is at most 1.

  - in the case of a dose response experiment, we also enforce that:
    * the difference in the total number of concentrations of a given compound between any two rows is at most 1.
    * the difference in the total number of concentrations of a given compound between any two columns is at most 1.
    * the difference in the total number of concentrations of a given compound between the upper half and the lower half of the microplate is at most 1.
    * the difference in the total number of concentrations of a given compound between the left half and the right half of the microplate is at most 1.
    * if the microplates are large enough, then the different concentrations of the same compound are not placed next to each other.

- Across all microplates:
  - the difference in the total number of samples between any two rows is at most 1.
  - the difference in the total number of samples between any two columns is at most 1.

### Section 1: Declarative file format for microplate experiments

Example of the declarative file format used to specify microplate experiments in MiniZinc syntax. The source code can be found in file small-example.dzn at https://github.com/pharmbio/plaid.

```
%%% Small Example File %%%

%% Plate dimentions:
num_rows = 4; %% height
num_cols = 6; %% width

% Number of times a plate is divided horizontally (with horizontal lines).
% 1 corresponds to using the whole plate,
% 2 corresponds to dividing the plate into 2 halves, one on top of the other. Both cells have the same layout.
horizontal_cell_lines = 1;

% Number of times a plate is divided vertically (with vertical lines).
% 1 corresponds to using the whole plate,
% 2 corresponds to dividing the plate into 2 halves, left and right. Both cells have the same layout.
vertical_cell_lines = 1;

allow_empty_wells = false; % Used as validation
size_empty_edge = 1;

% Turning on/off some constraints
concentrations_on_different_rows = true;
concentrations_on_different_columns = true;

% Restriction: either replicates_on_different_plates or replicates_on_same_plate must be false (or both)
replicates_on_different_plates = true;
replicates_on_same_plate = false;


%%% Compounds %%%
compounds = 5; %% number of drugs/compounds
compound_names = ["comp␣1", "comp␣2", "comp␣3", "comp␣4", "comp␣5" ];

compound_replicates = [2,2,2,2,2];
compound_concentrations = [2,2,2,2,2];


compound_concentration_names = [|"c11","c12"|"c21","c22"|"c31","c32"|"c41","c42"|"c51","c52"|];

%%% Used for drawing layouts in LaTeX %%%
compound_concentration_indicators = ["",""];


%%% Controls %%%
num_controls = 4;
control_replicates = [6,2,2,2];
control_concentrations = [1,1,1,1];
control_names = ["pos","neg","blank","DMSO"];
control_concentration_names =  [|"100"|"100"|"100"|"100"|];
```

### Supplementary figures

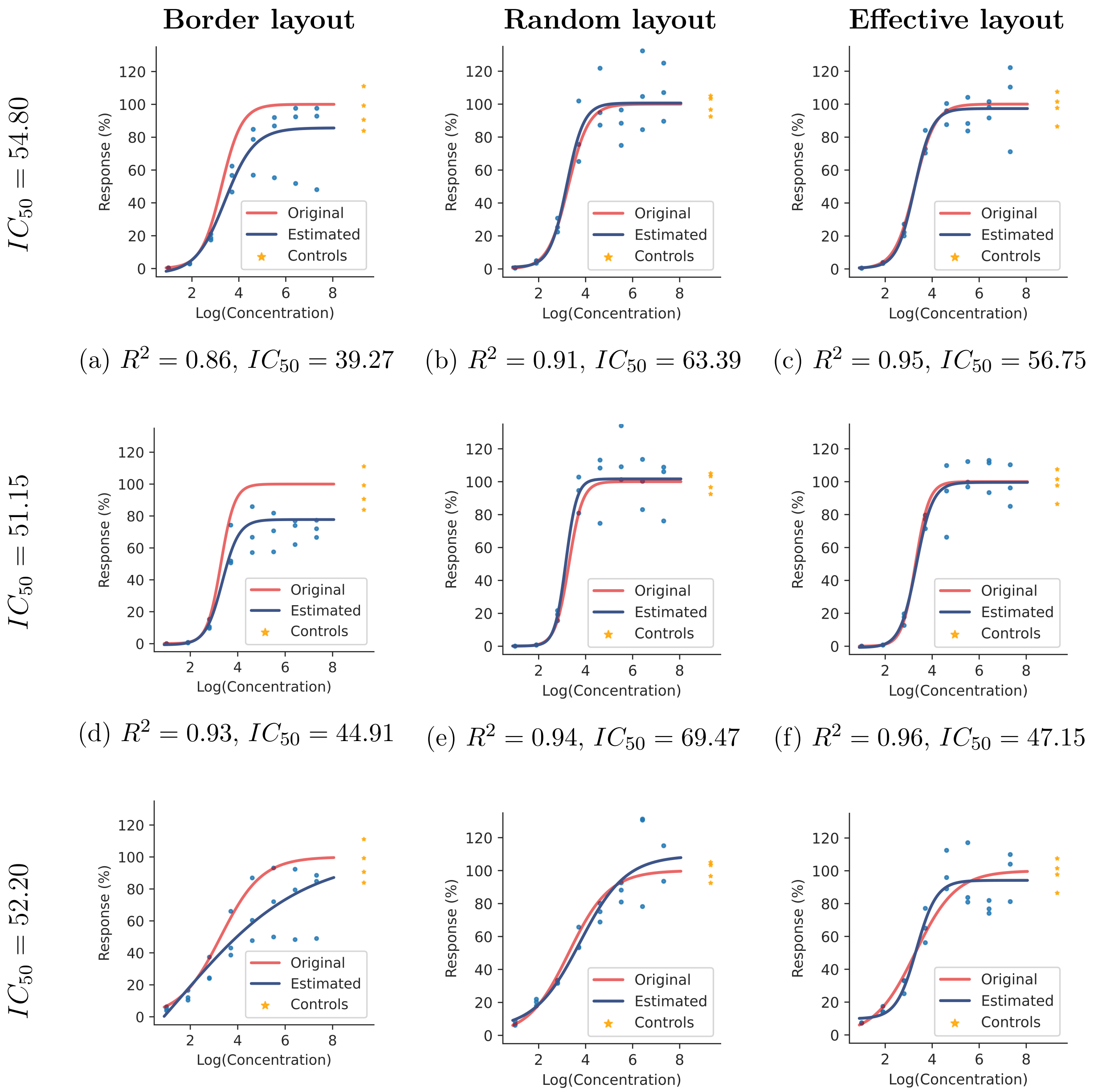


(a) $R^2 = 0.86$, $IC_{50} = 39.27$ (b) $R^2 = 0.91$, $IC_{50} = 63.39$ (c) $R^2 = 0.95$, $IC_{50} = 56.75$

(d) $R^2 = 0.93$, $IC_{50} = 44.91$ (e) $R^2 = 0.94$, $IC_{50} = 69.47$ (f) $R^2 = 0.96$, $IC_{50} = 47.15$

(g) $R^2 = 0.77$, $IC_{50} = 2896.72$ (h) $R^2 = 0.90$, $IC_{50} = 17.35$ (i) $R^2 = 0.91$, $IC_{50} = 46.81$

Figure 1: Examples of dose response curves for simulated experiments using 8 doses with 3 replicates (blue dots), 4 negative controls (yellow stars), and strong bowl-shaped plate effects. On the left of each row we show the expected $IC_{50}$ value. On the caption below each plot we show the obtained $R^2$ and $IC_{50}$ values.

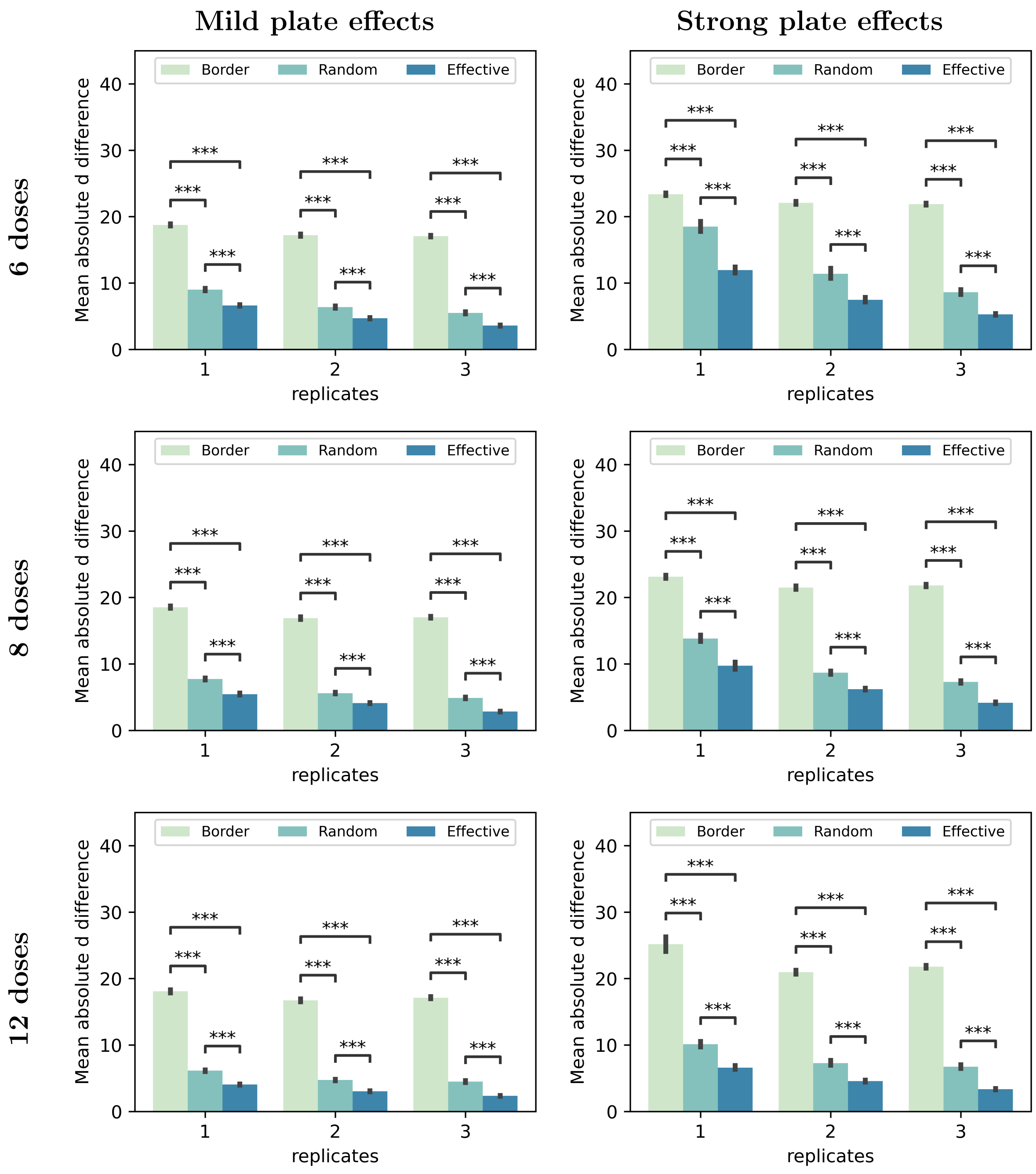


Figure 2: Mean absolute difference between expected and obtained heights (max value) for dose-response curves using various numbers of doses, replicates, and bowl-shaped plate-effect strengths. The 4PL sigmoide curves were fitted using only the compound data (not including negative controls). *** indicates $p \leq 10^{-43}$.

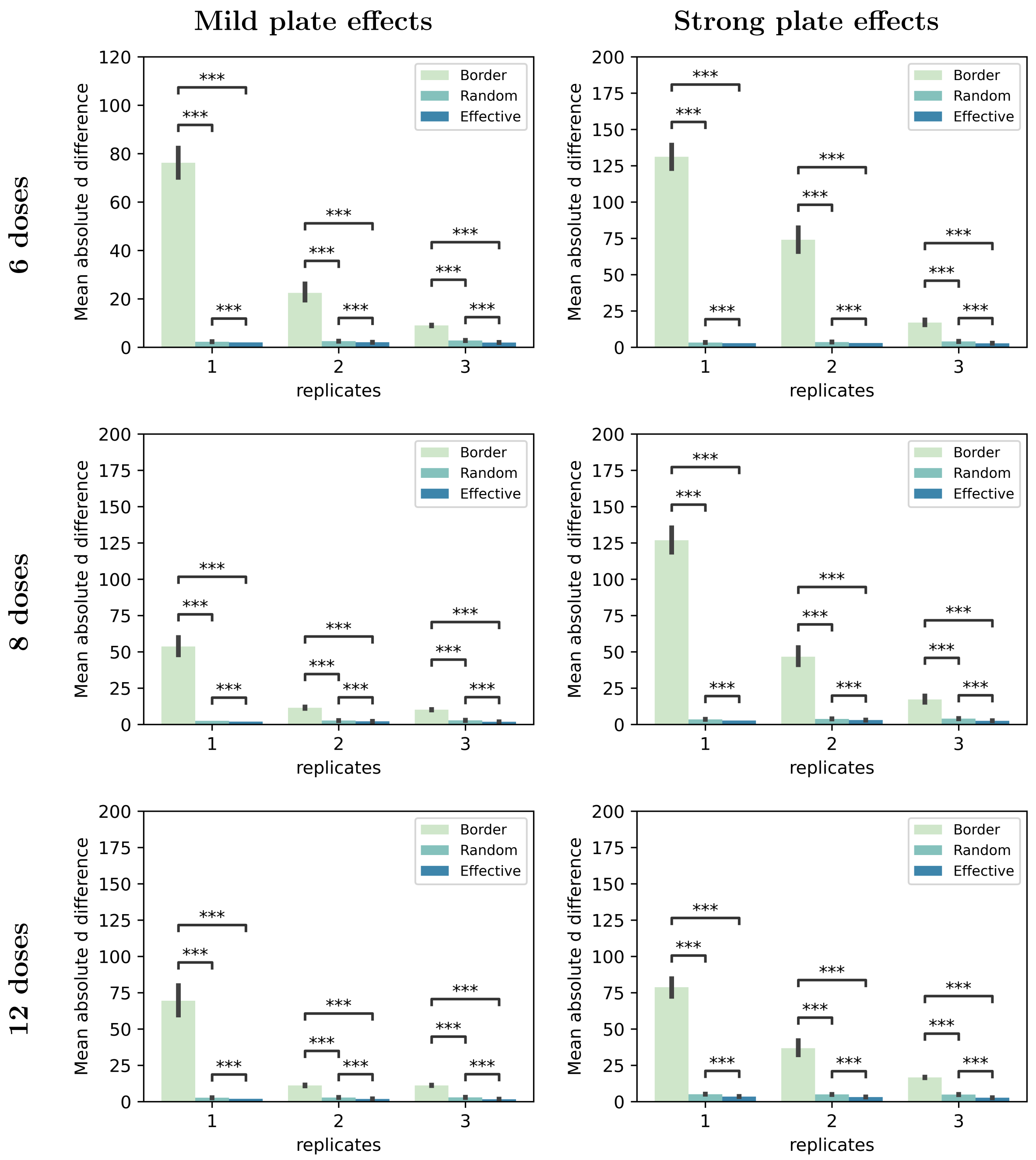


Figure 3: Mean absolute difference between expected and obtained heights (max value) for dose-response curves using various numbers of doses, replicates, and bowl-shaped plate-effect strengths. The 4PL sigmoide curves were fitted using compound data and 4 negative controls such that their mean was equal to the mean of the 20 negative controls the plate. *** indicates $p \leq 10^{-43}$.

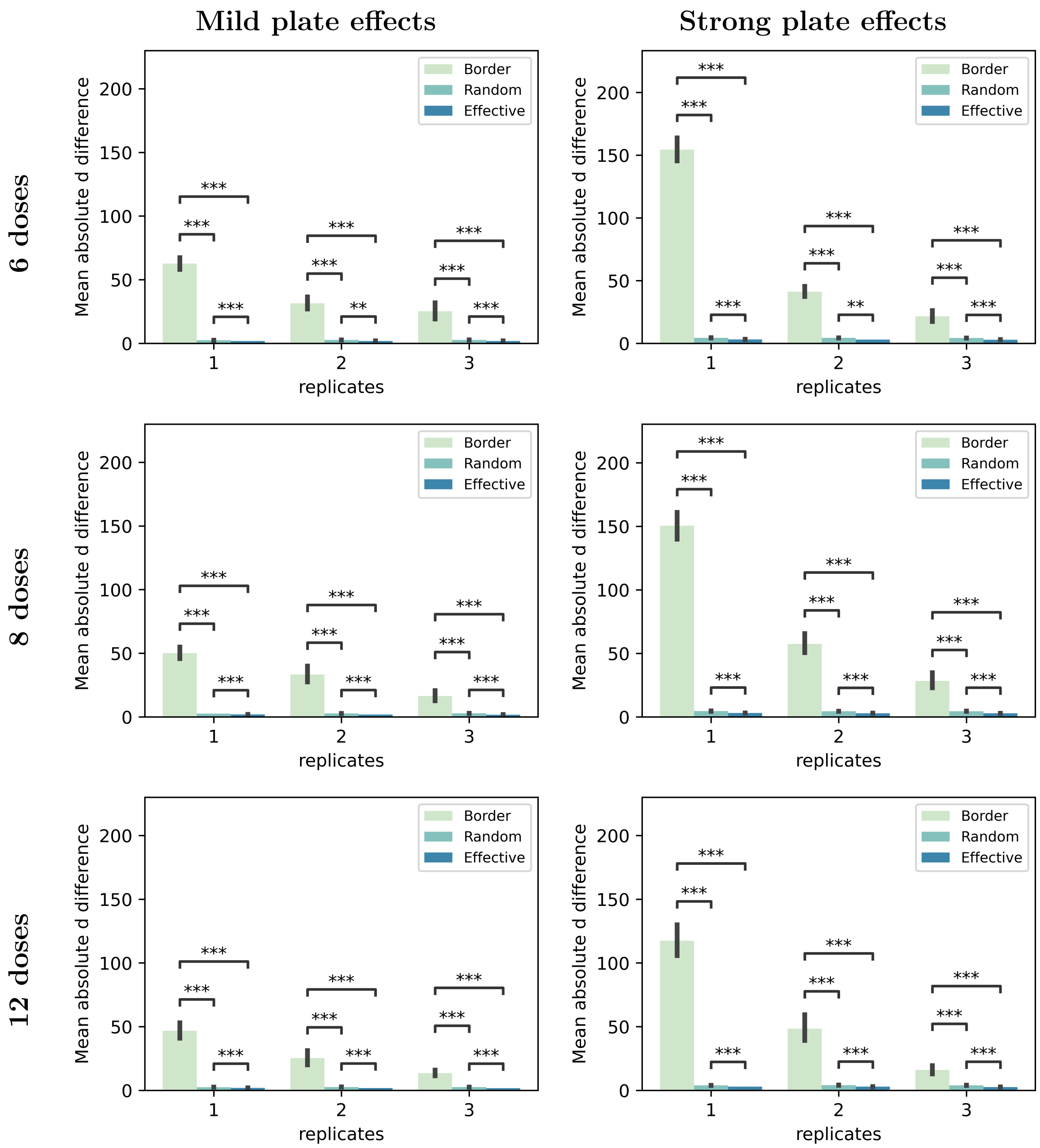


Figure 4: Mean absolute difference between expected and obtained heights (max value) for dose-response curves using various numbers of doses, replicates, and plate-effect strengths with a linear relationship to column number in the right half of the plate. The 4PL sigmoide curves were fitted using compound data and 4 negative controls such that their mean was equal to the mean of the 20 negative controls the plate. *** indicates $p \leq 10^{-43}$.

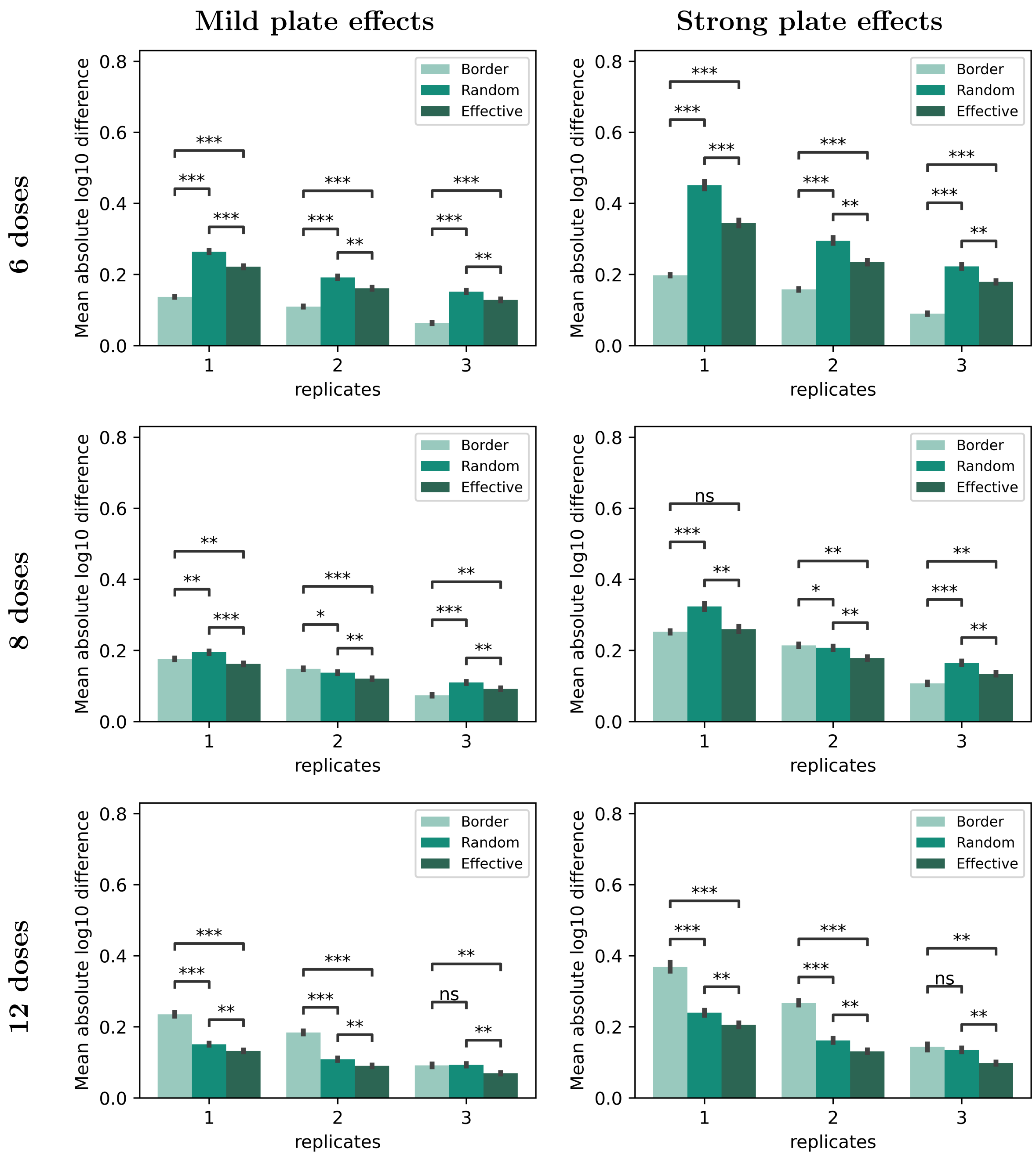


Figure 5: Mean absolute $\log_{10}$ difference between expected and obtained values of the relative $EC_{50}/IC_{50}$ for dose-response curves using various numbers of doses, replicates, and bowl-shaped plate-effect strengths. The 4PL sigmoide curves were fitted using only compound data. * indicates $p \leq 0.05$, ** indicates $p \leq 10^{-12}$, *** indicates $p \leq 10^{-43}$.

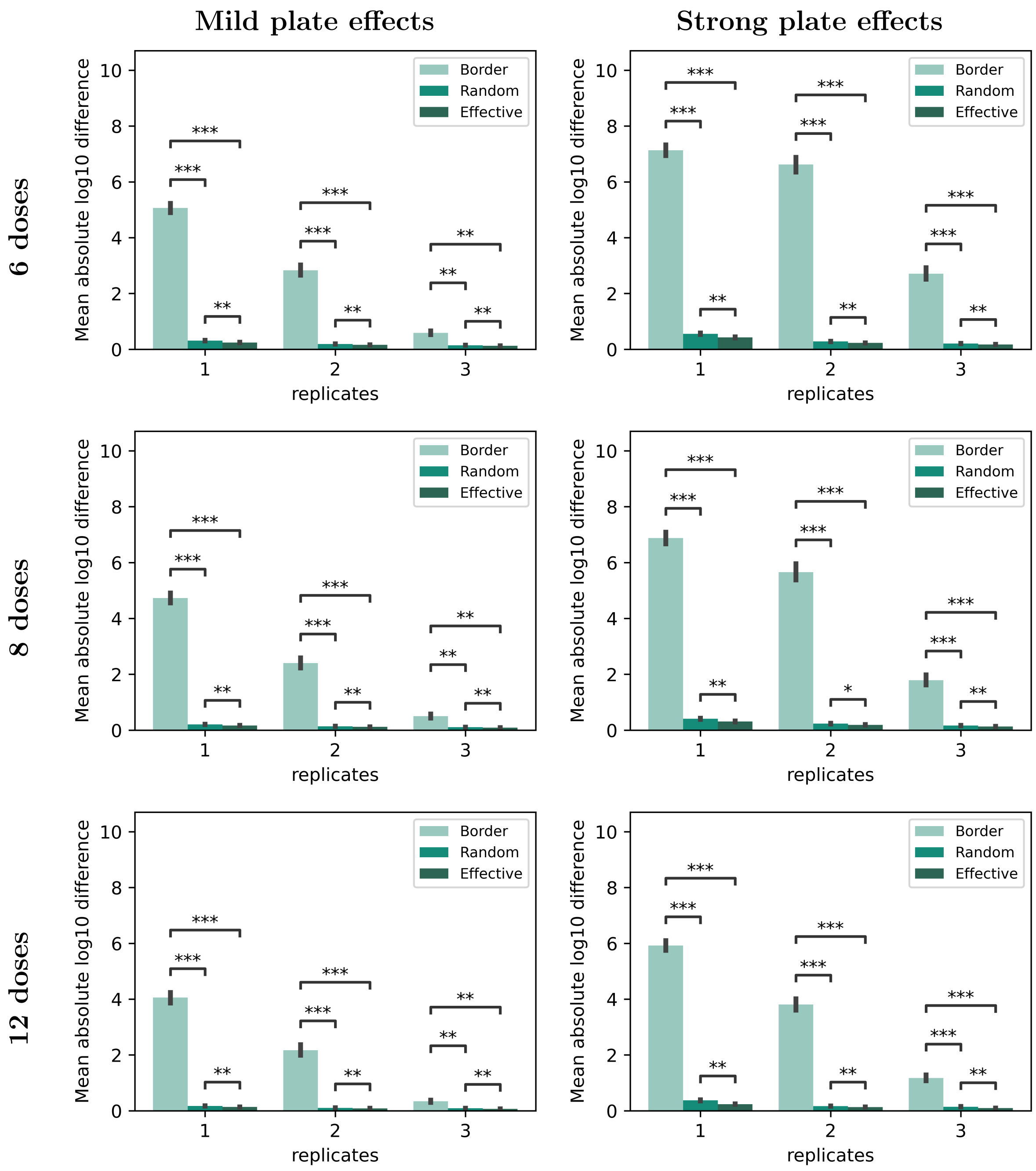


Figure 6: Mean absolute $\log_{10}$ difference between expected and obtained values of the relative $EC_{50}/IC_{50}$ for dose-response curves using various numbers of doses, replicates, and bowl-shaped plate-effect strengths. The 4PL sigmoide curves were fitted using compound data and 4 negative controls such that their mean was equal to the mean of the 20 negative controls the plate. * indicates $p \leq 10^{-4}$, ** indicates $p \leq 10^{-12}$, *** indicates $p \leq 10^{-43}$.

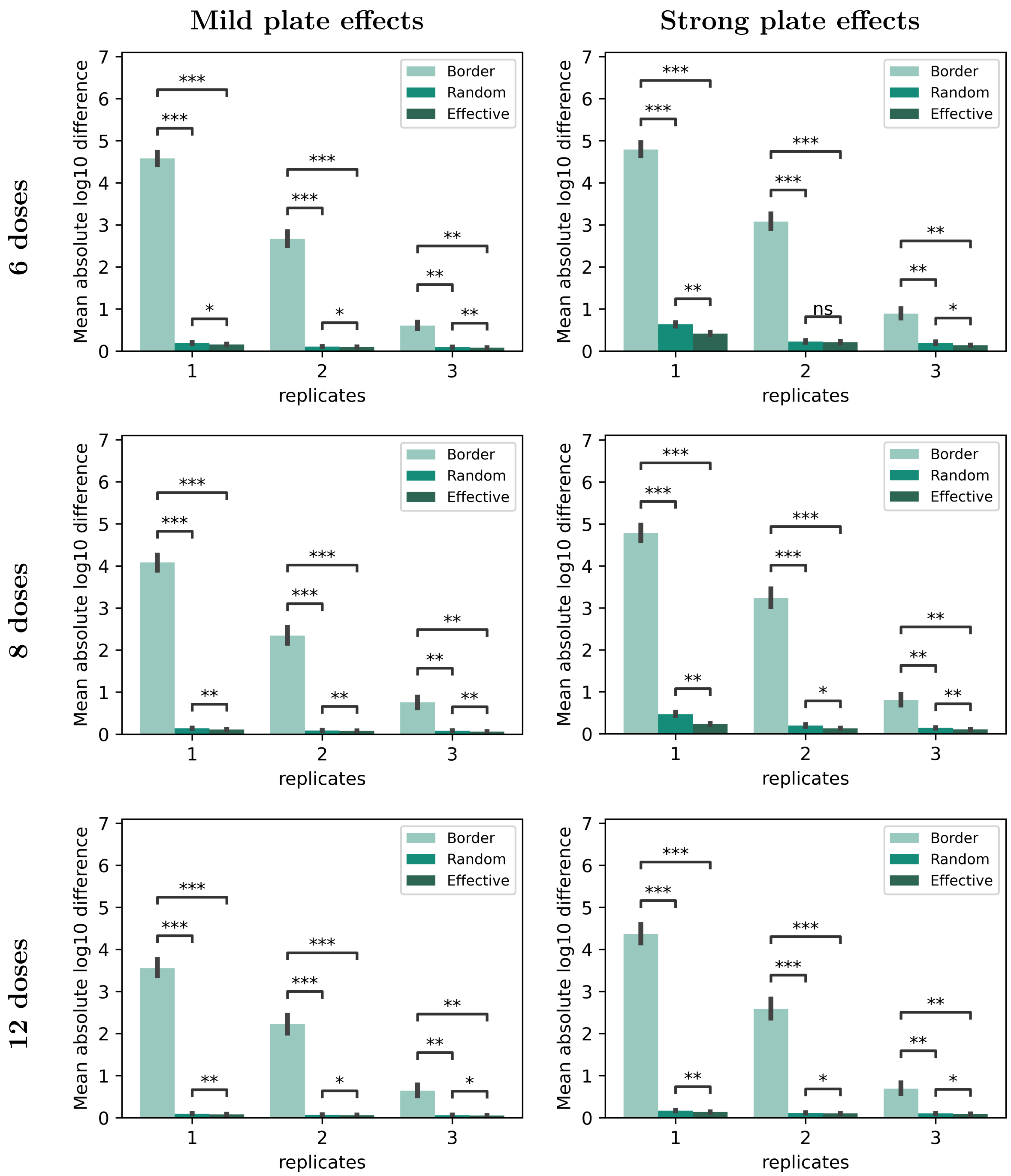


Figure 7: Mean absolute $\log_{10}$ difference between expected and obtained values of the relative $EC_{50}/IC_{50}$ for dose-response curves using various numbers of doses, replicates, and plate-effect strengths with a linear relationship to column number in the right half of the plate. The 4PL sigmoide curves were fitted using compound data and 4 negative controls such that their mean was equal to the mean of the 20 negative controls the plate. * indicates $p \leq 10^{-4}$, ** indicates $p \leq 10^{-12}$, *** indicates $p \leq 10^{-43}$.

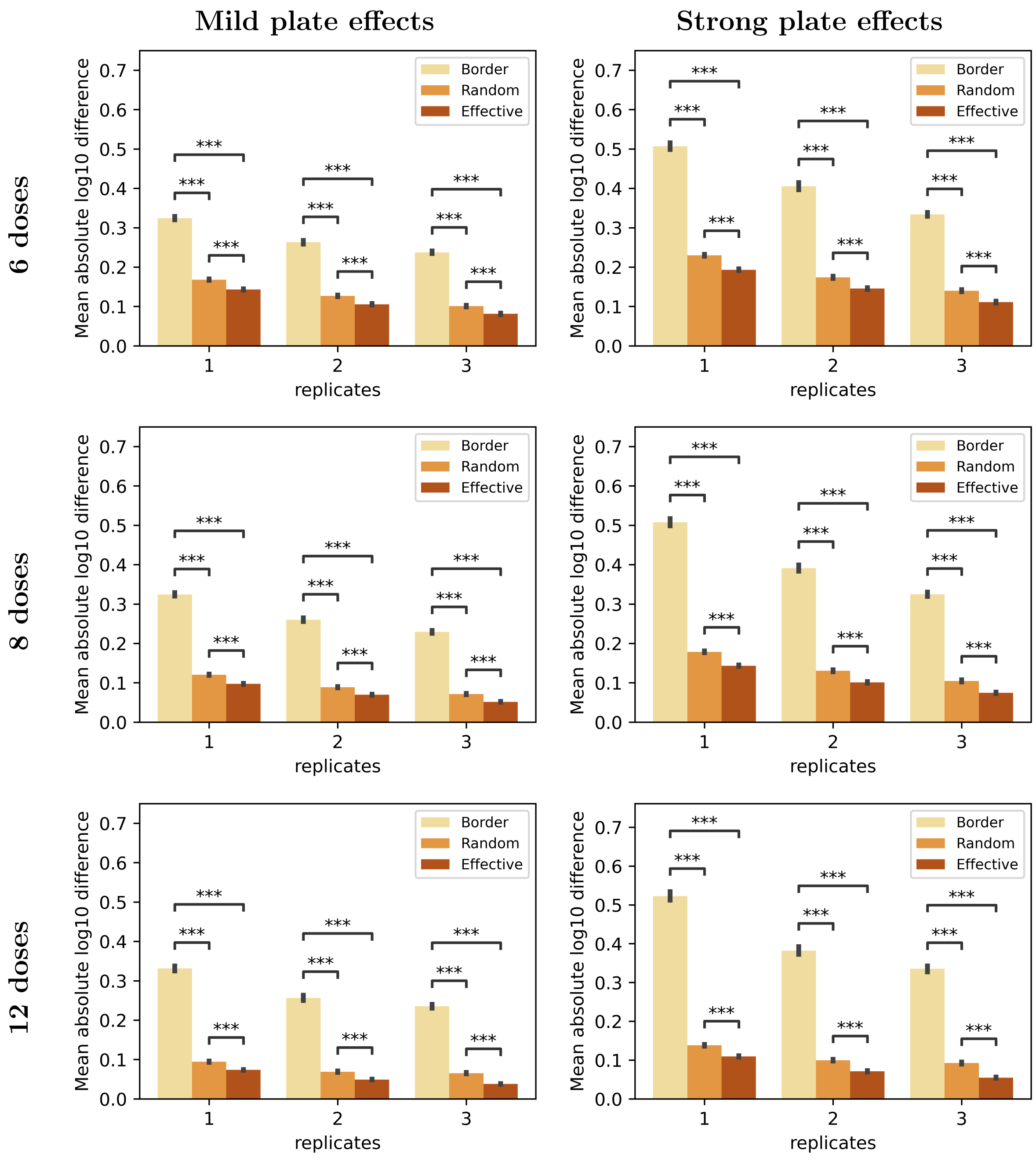


Figure 8: Mean absolute $\log_{10}$ difference between expected and obtained values of the absolute $EC_{50}/IC_{50}$ for dose-response curves using various numbers of doses, replicates, and bowl-shaped plate-effect strengths. The 4PL sigmoide curves were fitted using only compound data. *** indicates $p < 10^{-43}$.

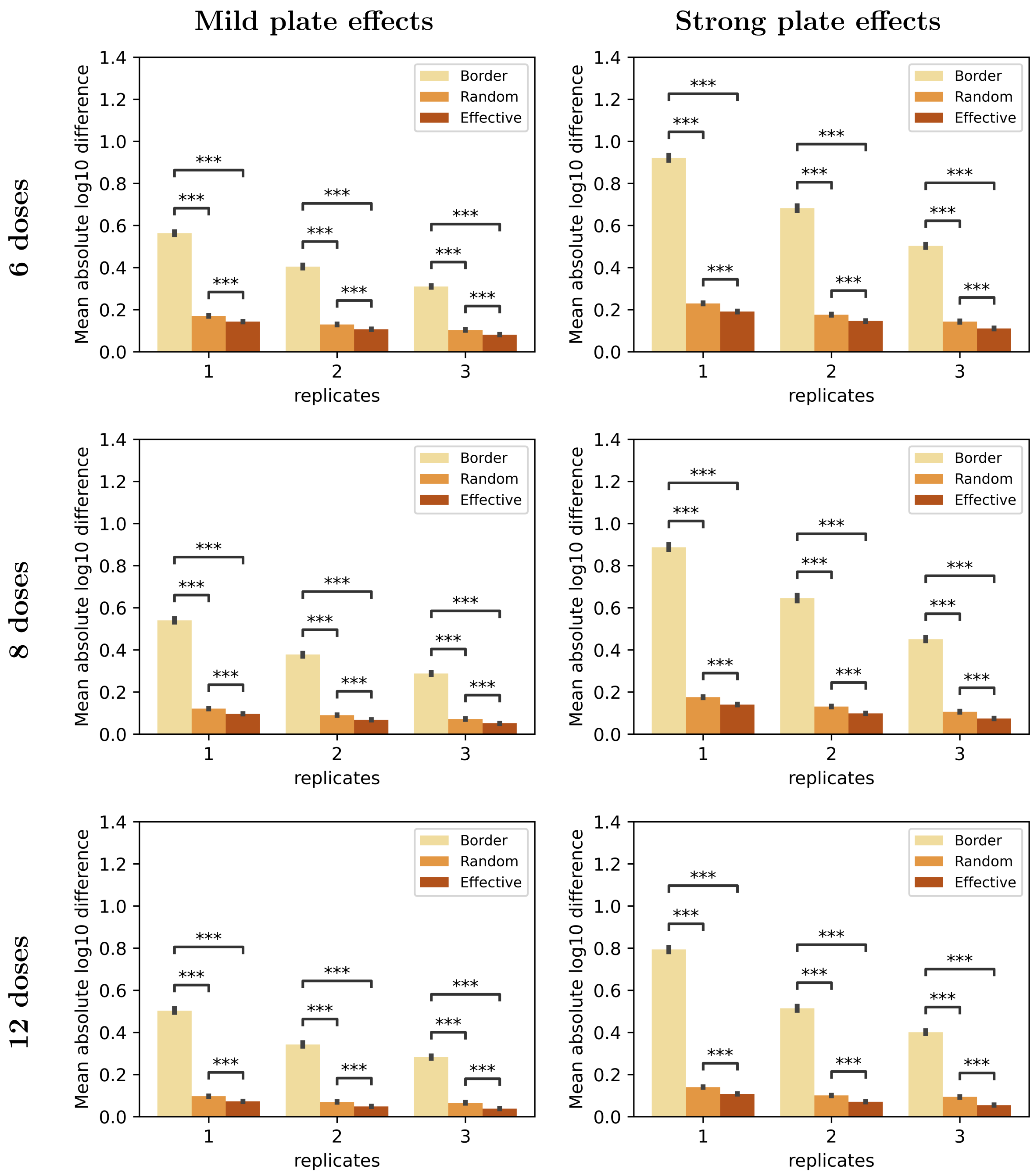


Figure 9: Mean absolute $\log_{10}$ difference between expected and obtained values of the absolute $EC_{50}/IC_{50}$ for dose-response curves using various numbers of doses, replicates, and bowl-shaped plate-effect strengths. The 4PL sigmoide curves were fitted using compound data and 4 negative controls such that their mean was equal to the mean of the 20 negative controls the plate. *** indicates $p \leq 10^{-43}$.

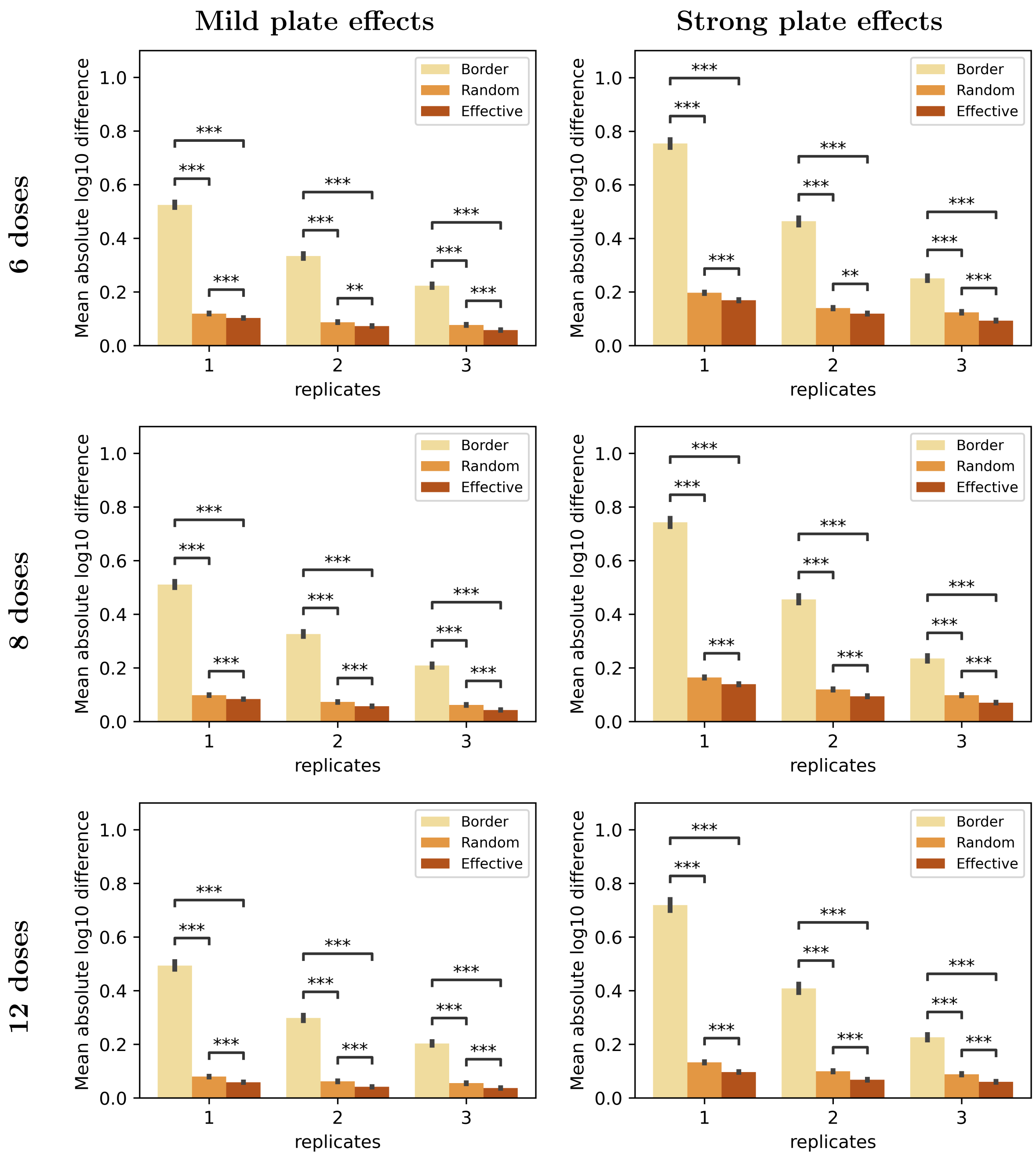


Figure 10: Mean absolute $\log_{10}$ difference between expected and obtained values of the absolute $EC_{50}/IC_{50}$ for dose-response curves using various numbers of doses, replicates, and plate-effect strengths with a linear relationship to column number in the right half of the plate. The 4PL sigmoide curves were fitted using compound data and 4 negative controls such that their mean was equal to the mean of the 20 negative controls the plate. *** indicates $p \leq 10^{-43}$.

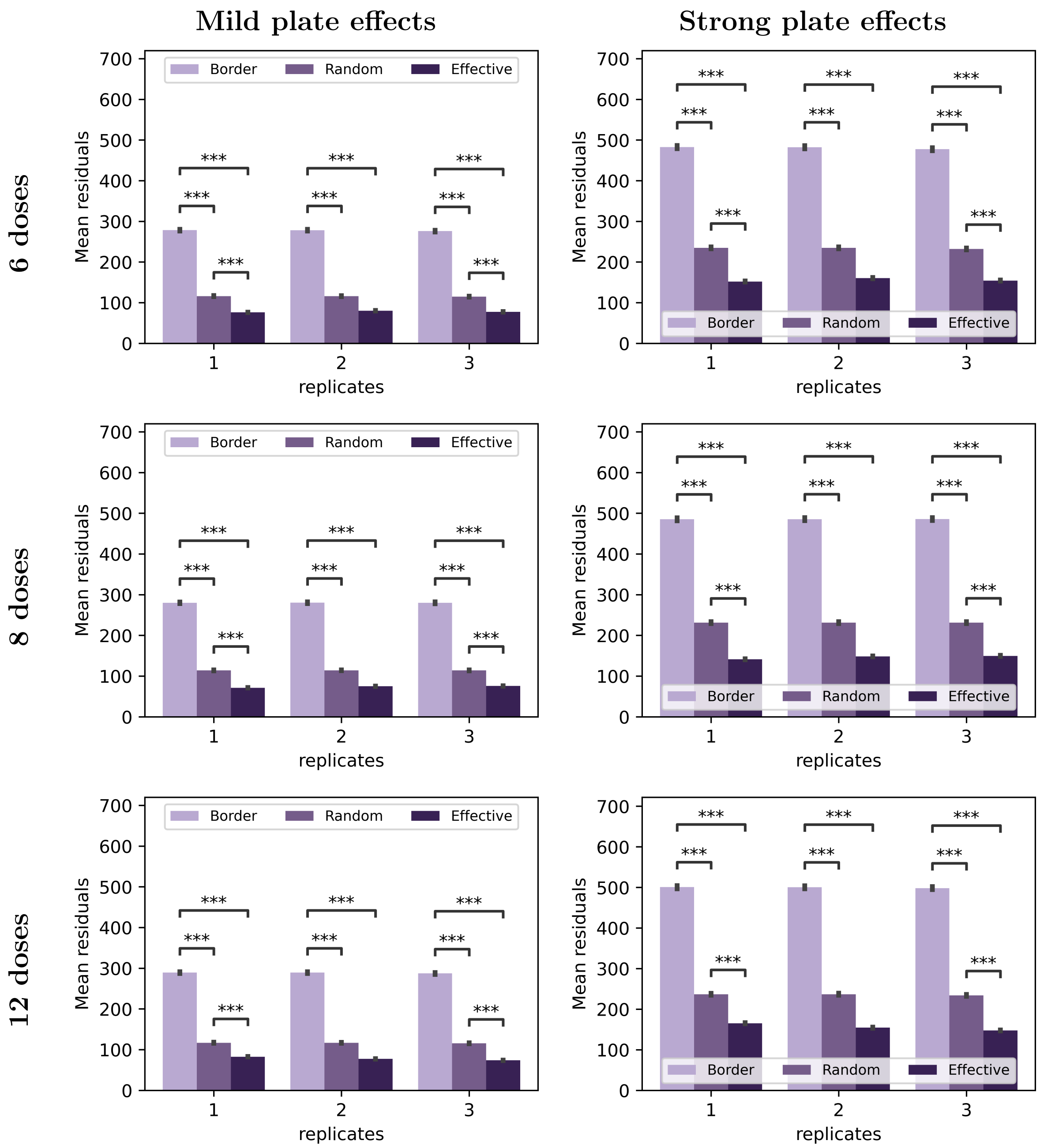


Figure 11: Residuals (MSE between expected and obtained responses) for dose-response simulations using various numbers of doses, replicates, and bowl-shaped plate-effect strengths. *** indicates $p \leq 10^{-43}$.

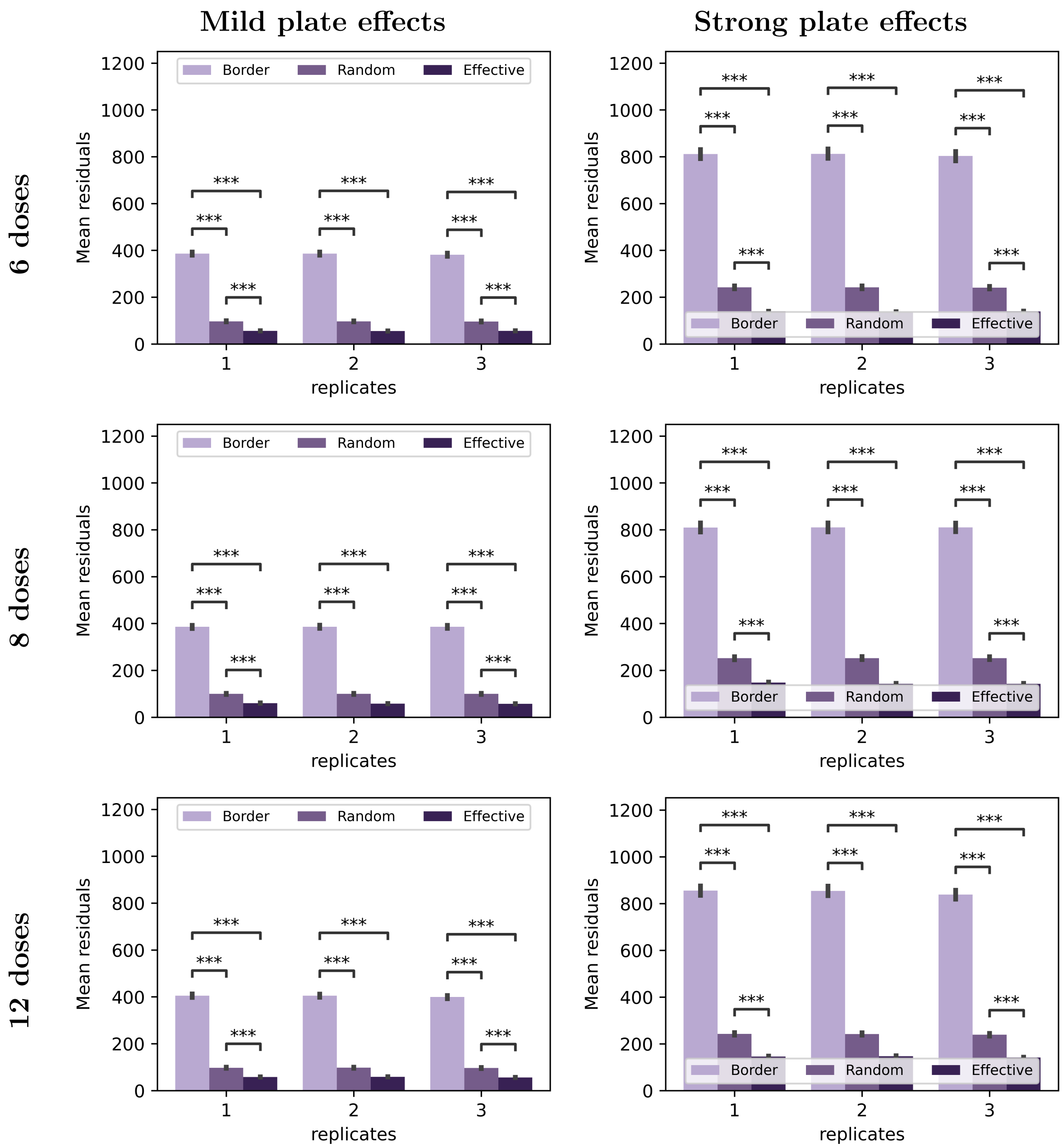


Figure 12: Residuals (MSE between expected and obtained responses) for dose-response simulations using various numbers of doses, replicates, and plate-effect strengths with a linear relationship to column number in the right half of the plate. *** indicates $p \leq 10^{-43}$.

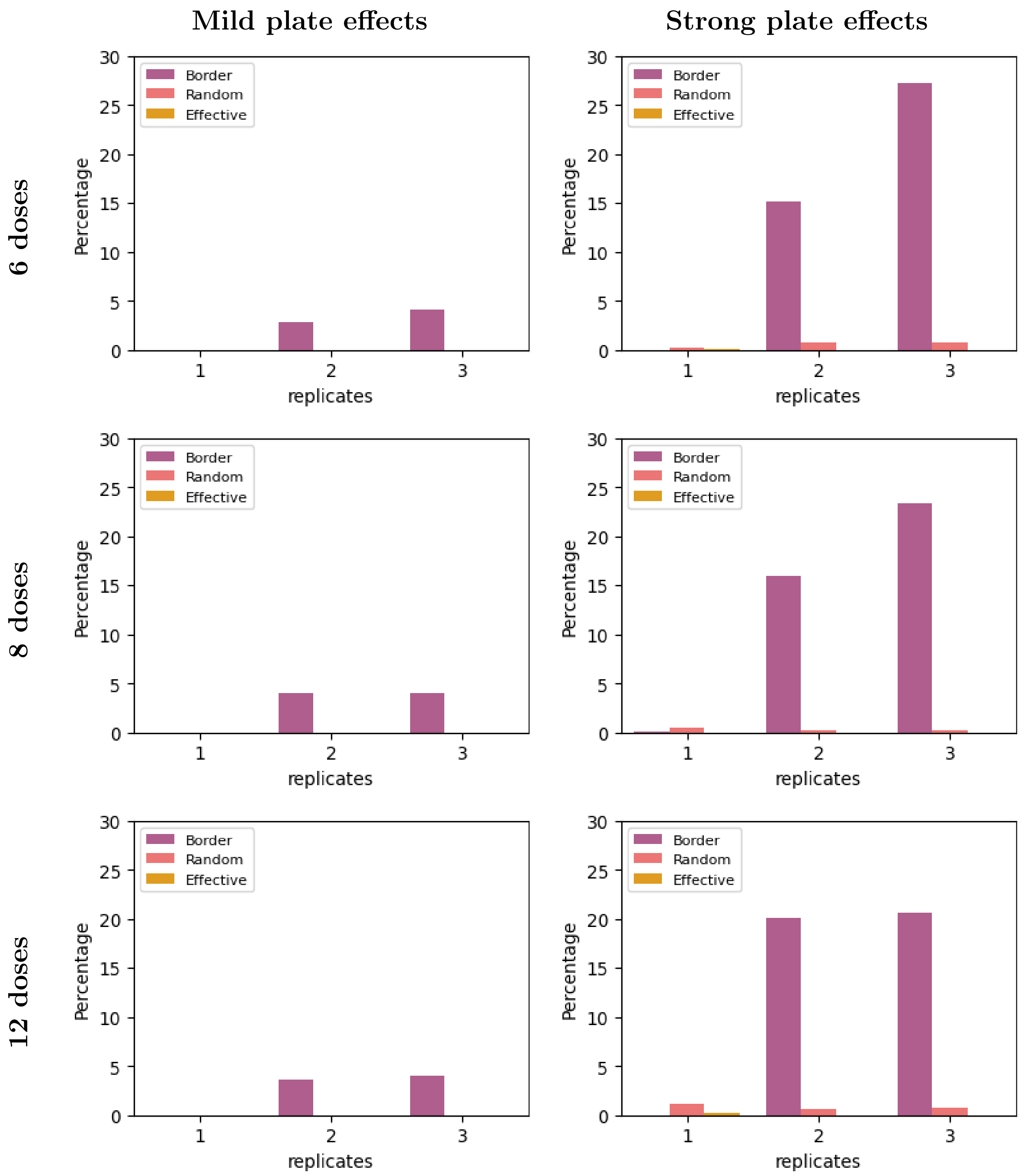


Figure 13: Percentage of dose-response curves with low-quality ($R^2 < 80\%$) using various numbers of doses, replicates, and bowl-shaped plate-effect strengths. The 4PL sigmoide curves were fitted using only compound data.

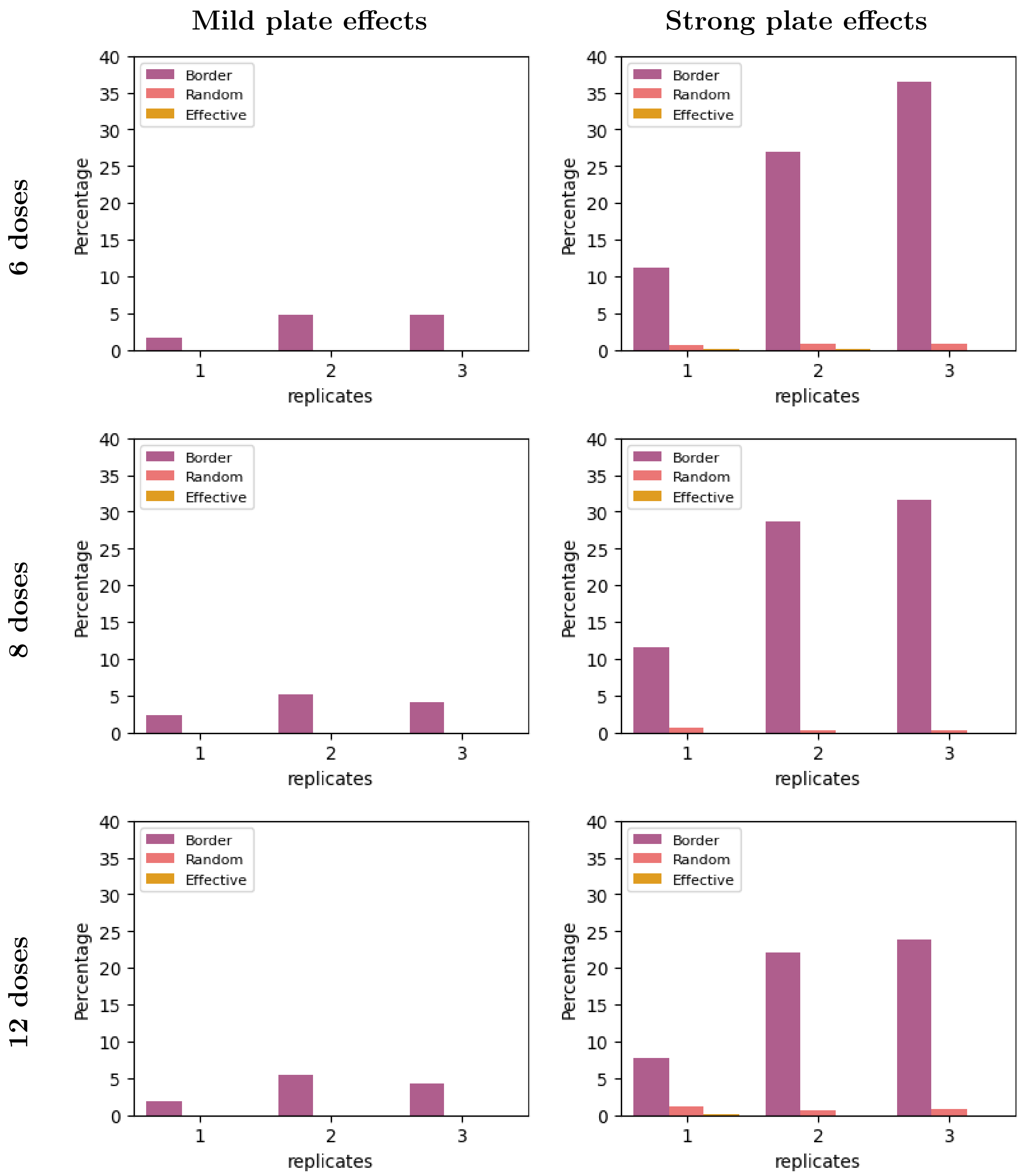


Figure 14: Percentage of dose-response curves with low-quality ($R^2 < 80\%$) using various numbers of doses, replicates, and bowl-shaped plate-effect strengths. The 4PL sigmoide curves were fitted using compound data and 4 negative controls such that their mean was equal to the mean of the 20 negative controls the plate.

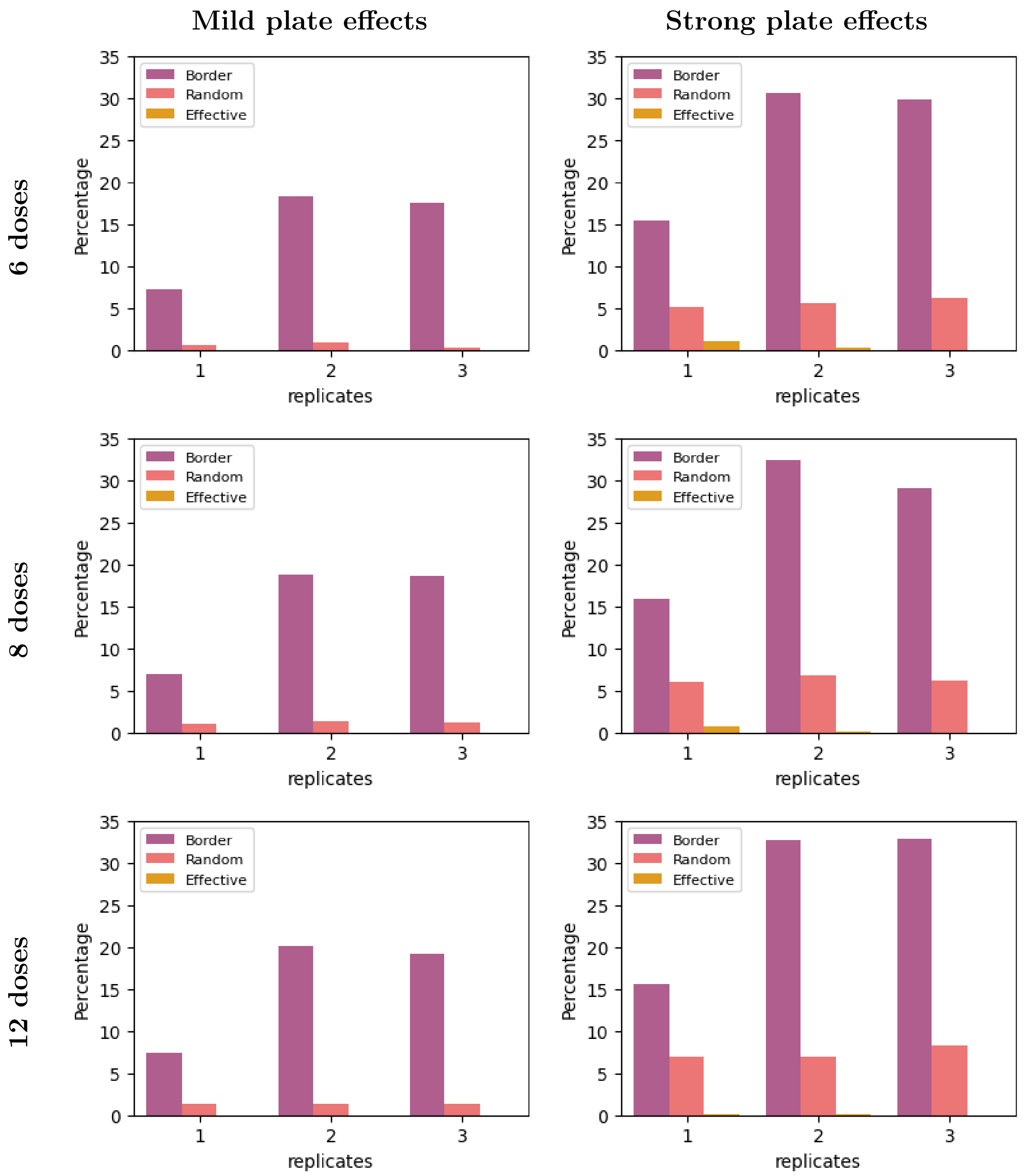


Figure 15: Percentage of dose-response curves with low-quality ($R^2 < 80\%$) using various numbers of doses, replicates, and plate-effect strengths with a linear relationship to column number in the right half of the plate. The 4PL sigmoide curves were fitted using compound data and 4 negative controls such that their mean was equal to the mean of the 20 negative controls the plate.

(a) Border (b) Random (c) Effective

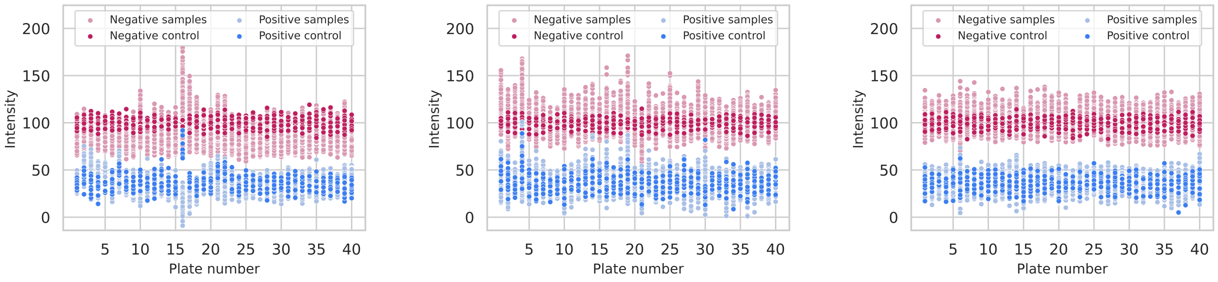


Figure 16: Comparison between expected and obtained values for screenings experiments using 10 positive and 10 negative controls on 384-well plate with only 1 replicate, 33% hit-rate and a mild bowl-shaped plate effects.

(a) Border (b) Random (c) Effective

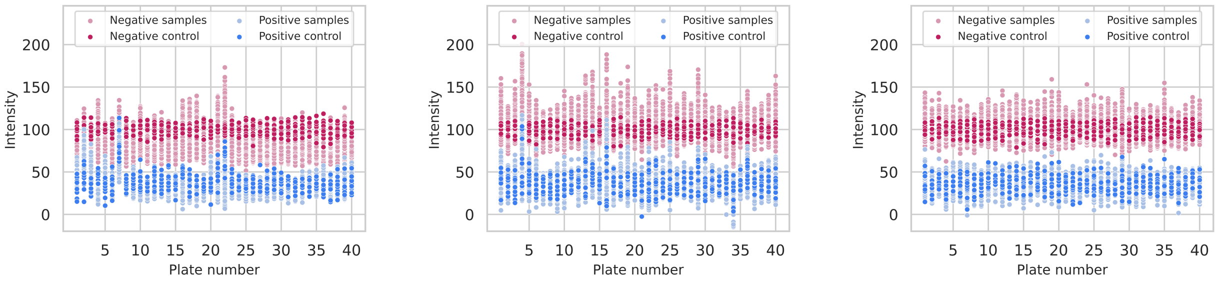


Figure 17: Comparison between expected and obtained values for screenings experiments using 10 positive and 10 negative controls on 384-well plate with only 1 replicate, 33% hit-rate and a mild-moderate bowl-shaped plate effects.

(a) Border (b) Random (c) Effective

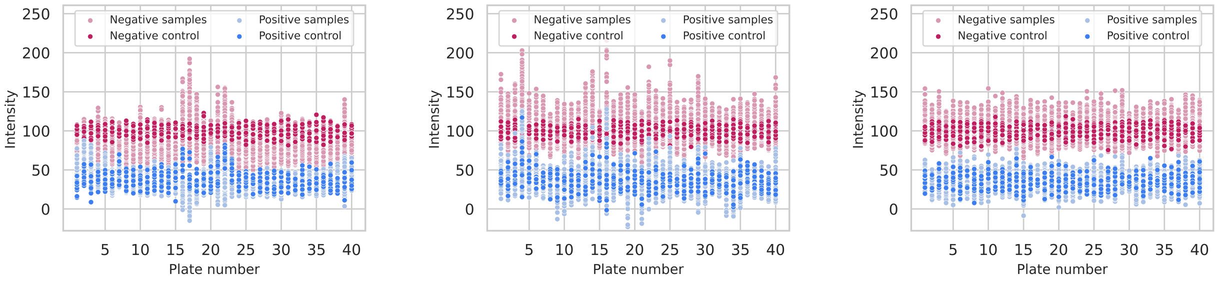


Figure 18: Comparison between expected and obtained values for screenings experiments using 10 positive and 10 negative controls on 384-well plate with only 1 replicate, 33% hit-rate and moderate bowl-shaped plate effects.

(a) Border (b) Random (c) Effective

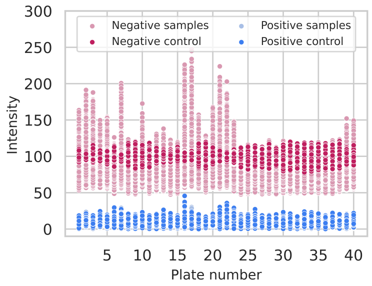


Figure 19: Comparison between expected and obtained values for screening experiments using 10 positive and 10 negative controls on 384-well plates with only 1 replicate, 1% hit-rate and moderately-strong bowl-shaped plate effects.

(a) Border (b) Random (c) Effective

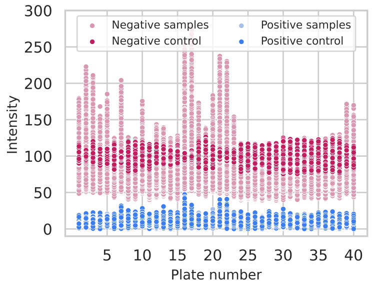


Figure 20: Comparison between expected and obtained values for screening experiments using 10 positive and 10 negative controls on 384-well plates with only 1 replicate, 1% hit-rate and strong bowl-shaped plate effects.

(a) Border (b) Random (c) Effective

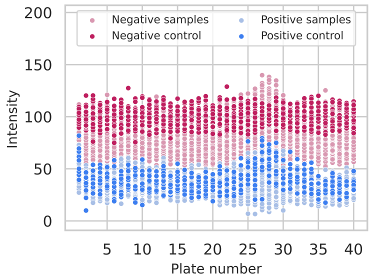


Figure 21: Comparison between expected and obtained values for screening experiments using 10 positive and 20 negative controls on 384-well plates with only 1 replicate, 33% hit-rate and moderate bowl-shaped plate effects.

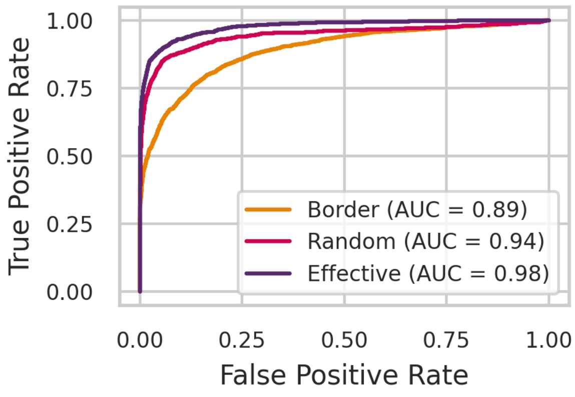


(a) 10% hits

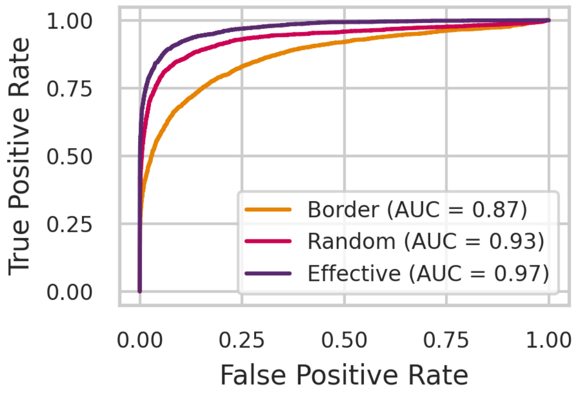


(b) 20% hits

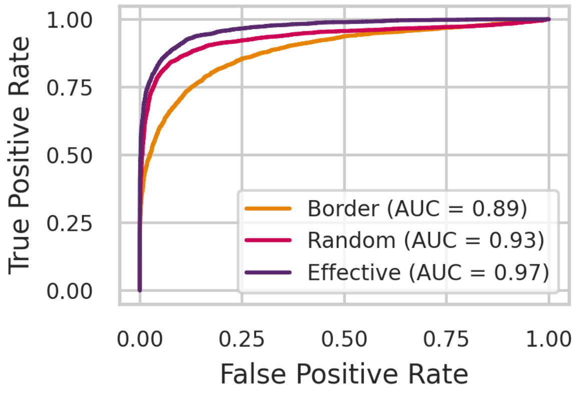


(c) 30% hits

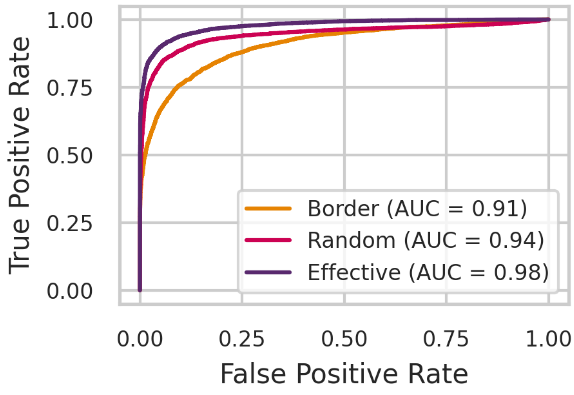


(d) 40% hits

Figure 22: Comparison of ROC curves for screening experiments with different hit rates, using layouts with 10 positive and 10 negative controls in the presence of strong bowl-shaped plate effects.

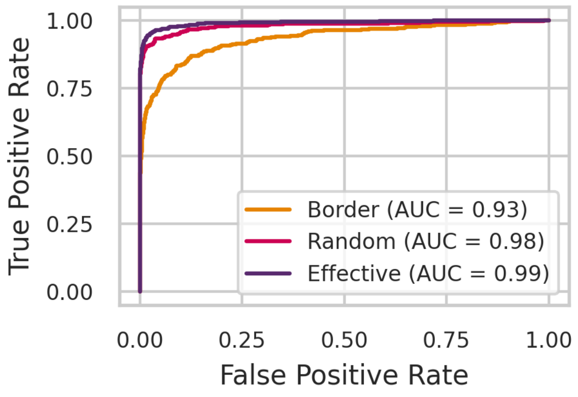


(a) 1% hits

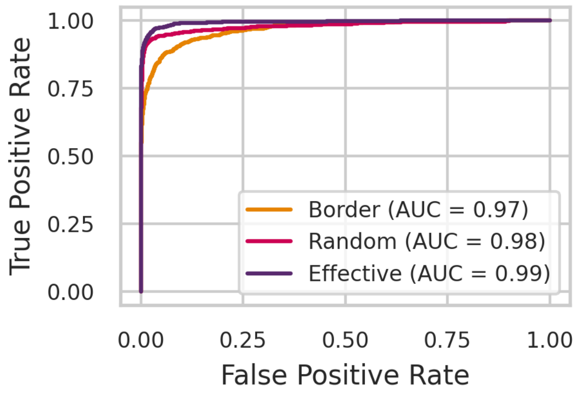


(b) 5% hits

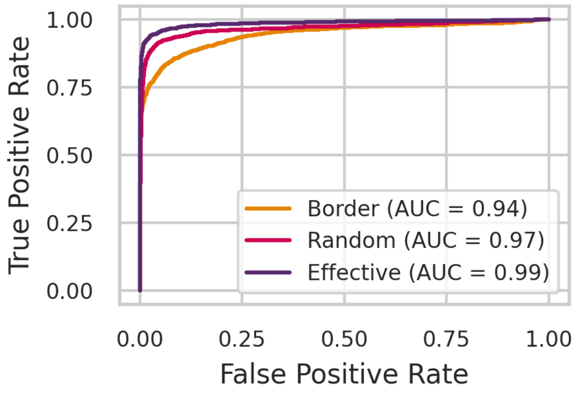


(c) 10% hits

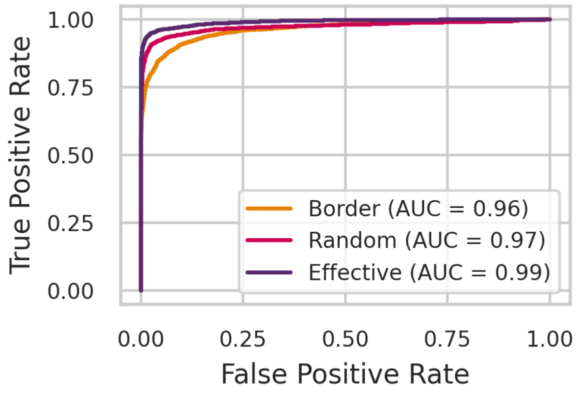


(d) 20% hits

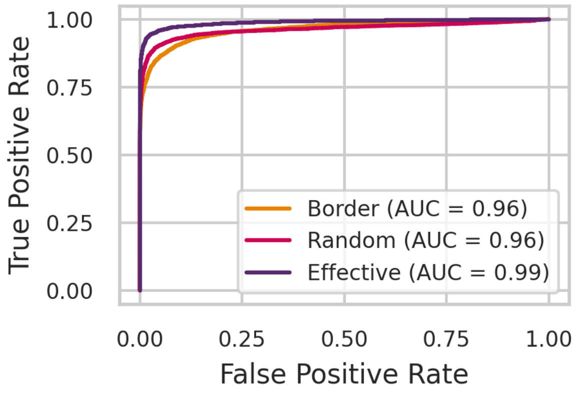


(e) 30% hits

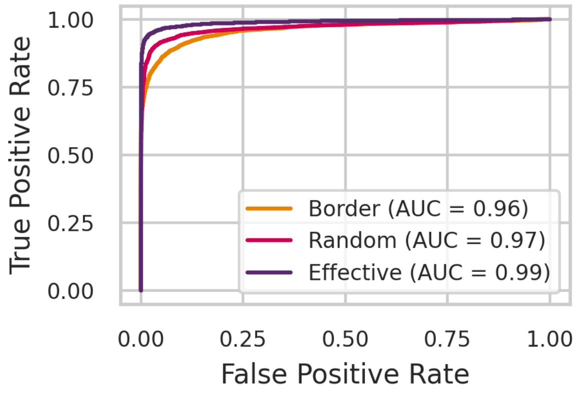


(f) 40% hits

Figure 23: Comparison of ROC curves for screening experiments with different hit rates, using layouts with 8 positive and 8 negative controls in the presence of strong bowl-shaped plate effects.

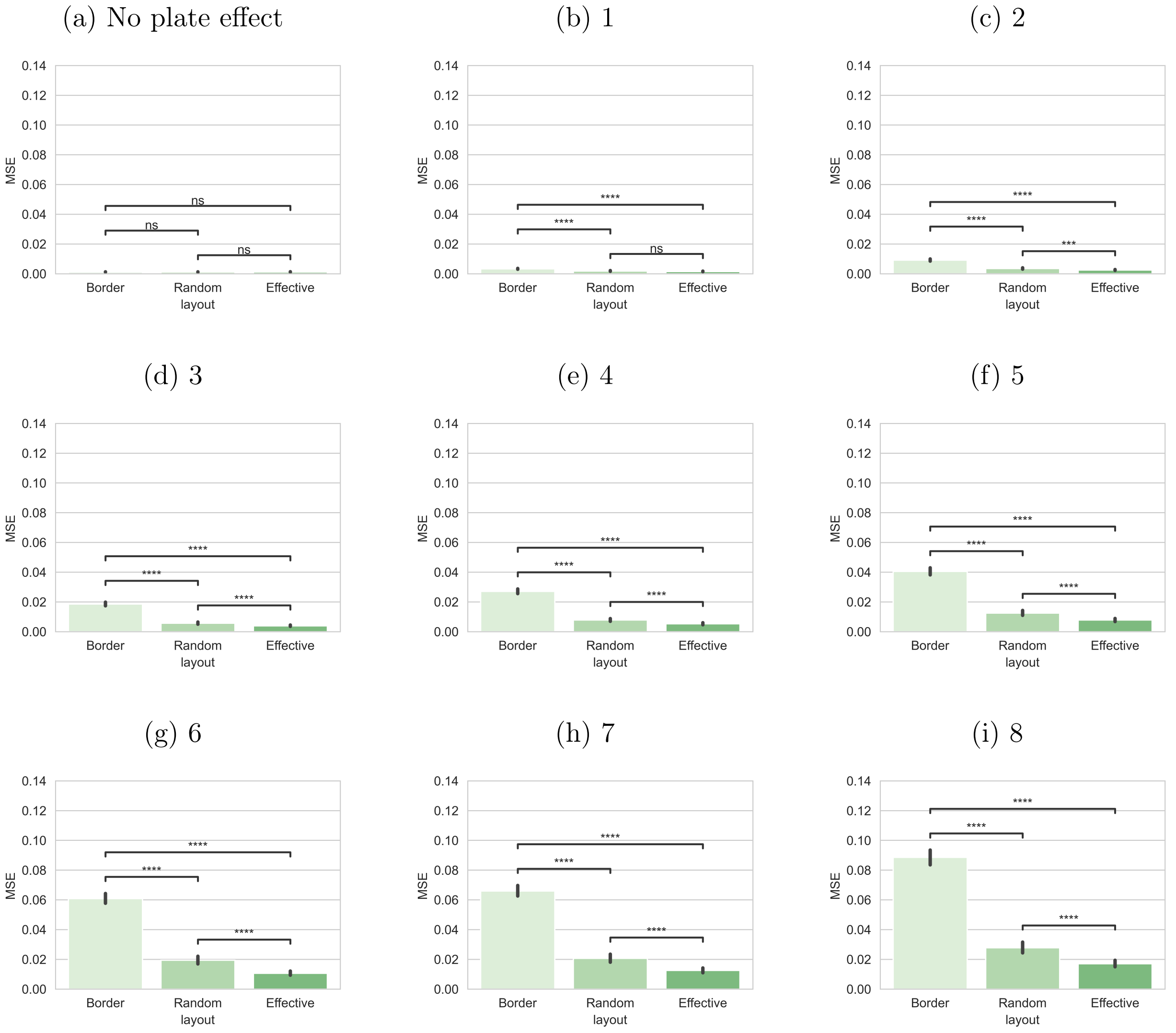


Figure 24: Comparison of MSE when calculating the Z' factor of plates for screenings experiments using 10 positive and 10 negative controls on 384-well plate with only 1 replicate, 1% hit-rate and increasing strengths of bowl-shaped plate effects. * indicates $p < 0.05$, ** indicates $p < 10^{-2}$, *** indicates $p < 10^{-3}$.

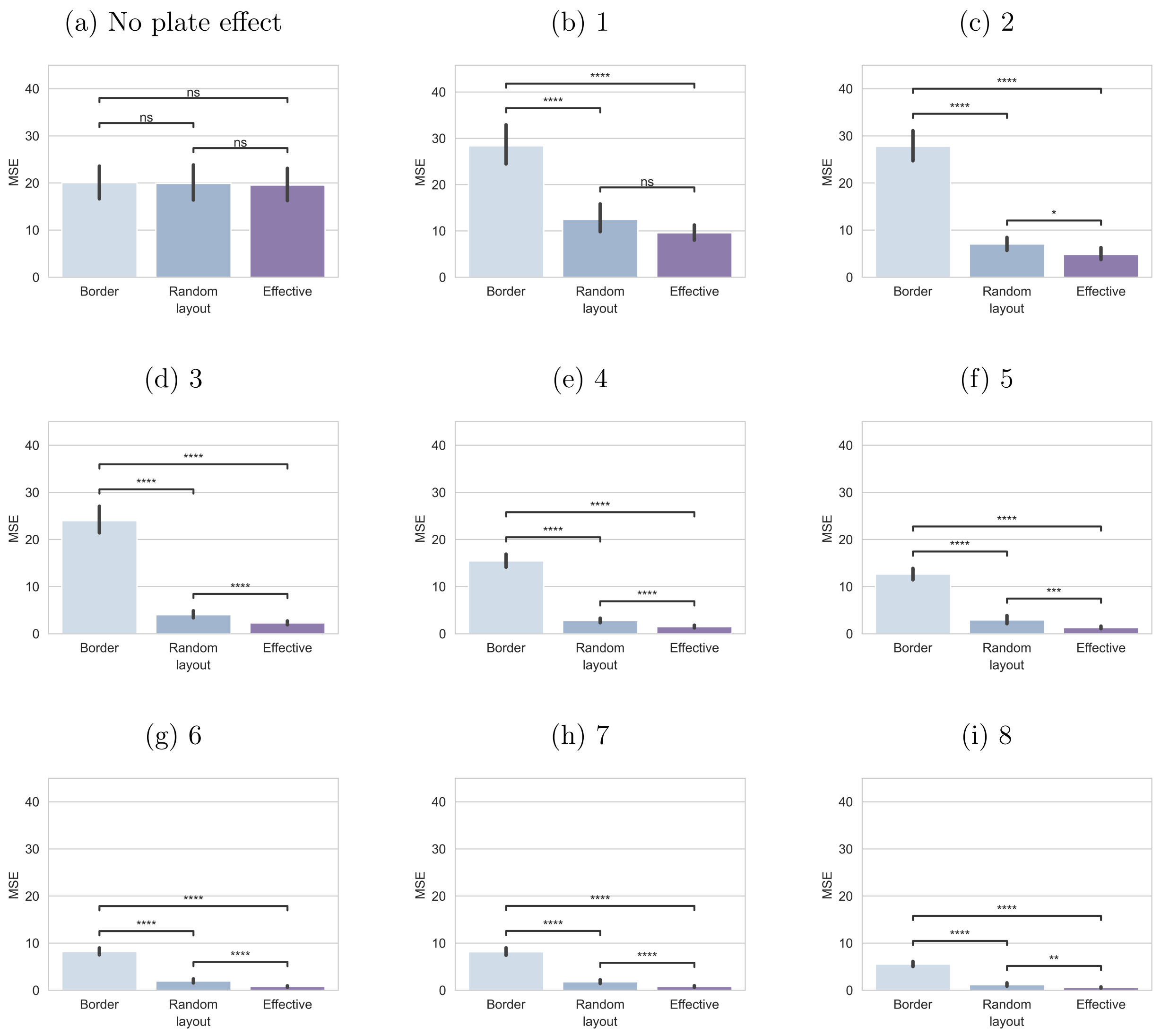


Figure 25: Comparison of MSE when calculating the SSMD of plates for screenings experiments using 10 positive and 10 negative controls on 384-well plate with only 1 replicate, 1% hit-rate and varying strengths of bowl-shaped plate effects. * indicates $p < 0.05$, ** indicates $p < 10^{-2}$, *** indicates $p < 10^{-3}$.

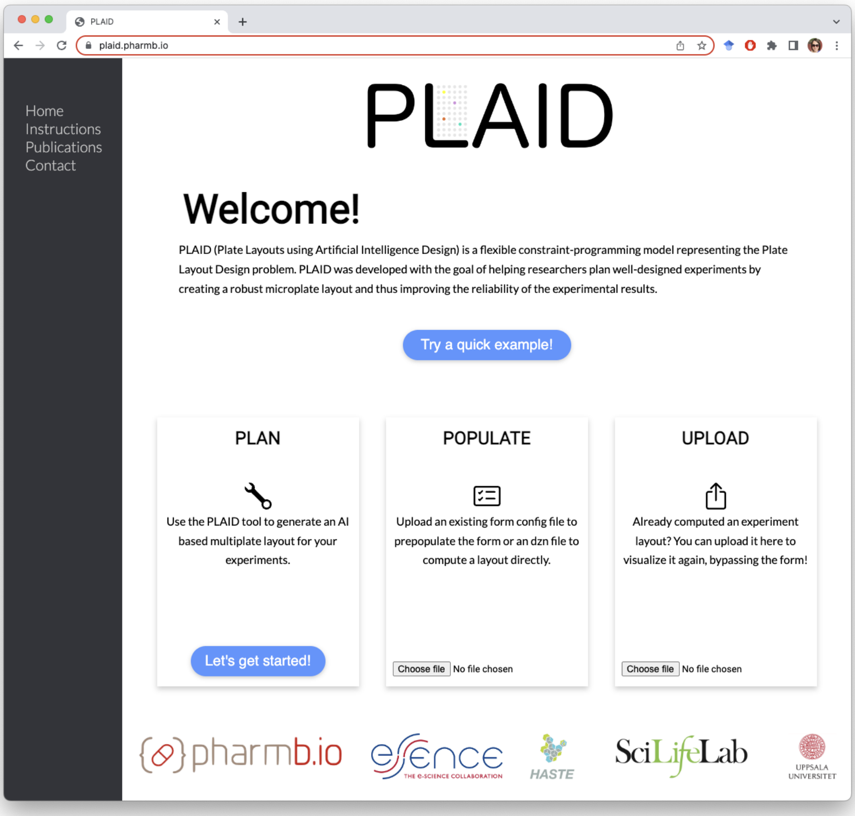


Figure 26: Homepage of the PLAID website, available at `plaid.pharmb.io`. Bottom-left: creating a new plan by typing in the experimental details, which can be saved in a .json format. Bottom-centre: import a file containing the experimental details using either our .dzn format or our .json format. Bottom-right: import a layout for visualisation. It only accepts layouts that have been downloaded in a .json format from the PLAID website.

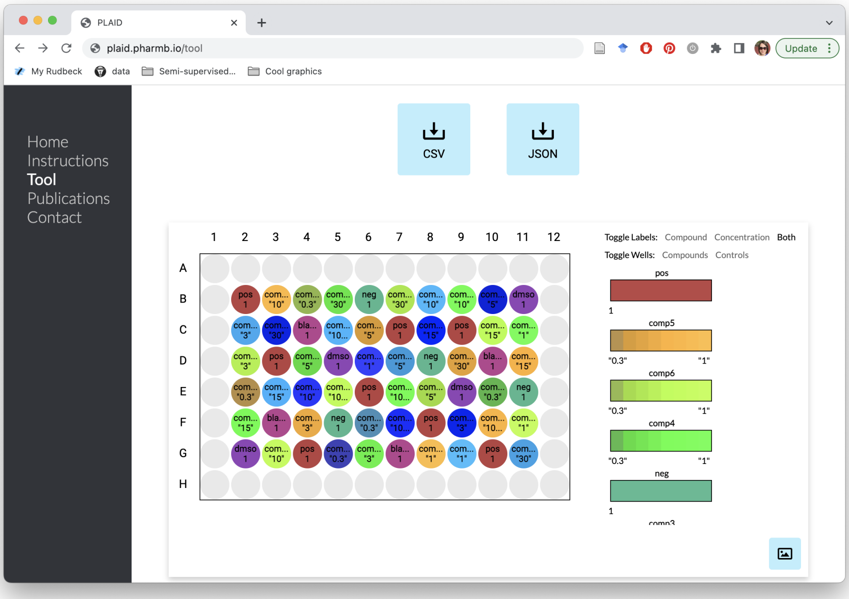


Figure 27: Layout of a 96-well plate as displayed on the PLAID website. This is one plate out of four that can be obtained for the experimental details in pl-example01.dzn (same as pl-example01.json). Both files are available at `https://github.com/pharmbio/plaid`.

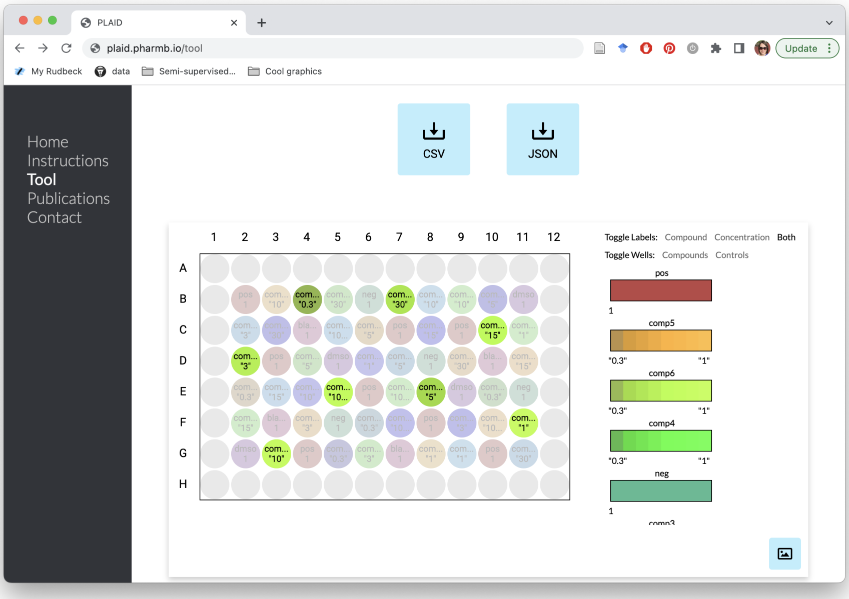


Figure 28: Layout of a 96-well plate as displayed on the PLAID website with comp6 being highlighted. Individual compounds and controls can be selected for easier visual inspection by clicking on their names in the list to right of the layout. This is one plate out of four that can be obtained for the experimental details in pl-example01.dzn (same as pl-example01.json). Both files are available at `https://github.com/pharmbio/plaid`.

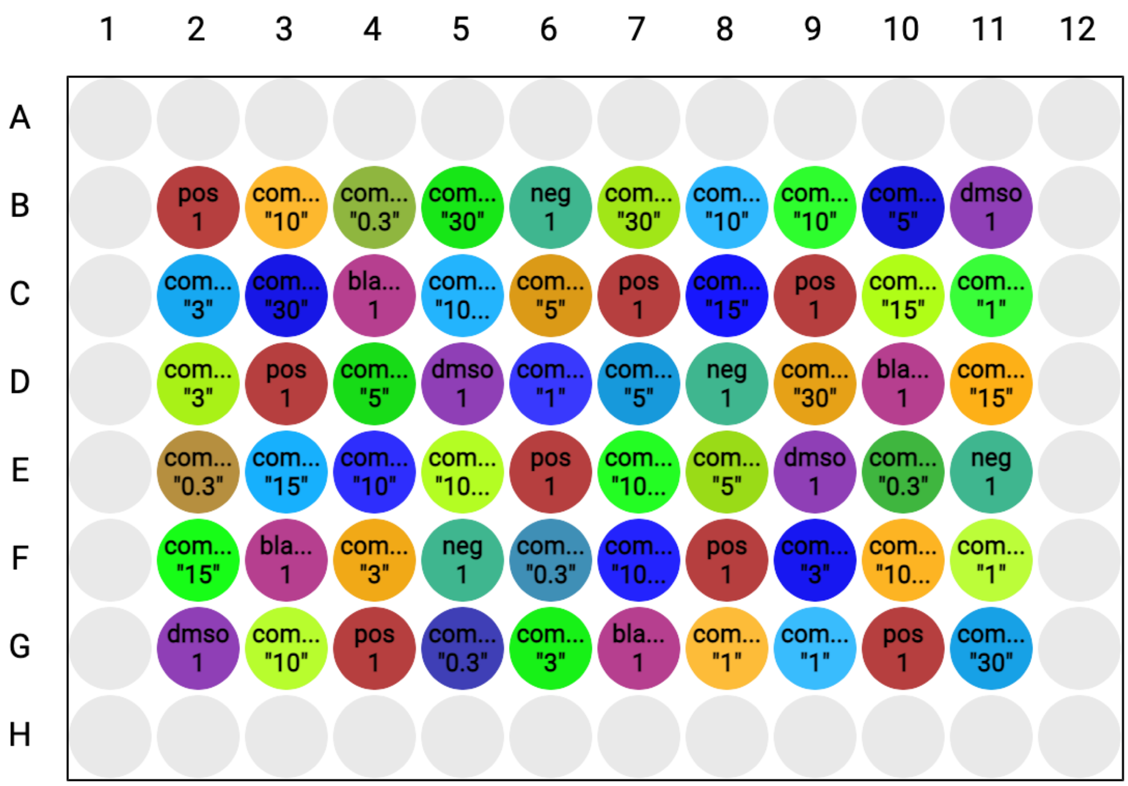


Figure 29: Layout of a 96-well plate as downloaded from the PLAID website. This is one plate out of four that can be obtained for the experimental details in pl-example01.dzn (same as pl-example01.json). Both files are available at `https://github.com/pharmbio/plaid`.

### Supplementary tables

| **Measurement** | **Comparison** | **1 replicate** | **2 replicates** | **3 replicates** |
|---|---|---|---|---|
| Relative $EC_{50}/IC_{50}$ | Effective – Random | 4.64e-25 | 1.21e-10 | 3.34e-28 |
| | Effective – Border | 0.00e+00 | 4.61e-172 | 4.30e-26 |
| | Random – Border | 0.00e+00 | 4.81e-164 | 1.47e-23 |
| Absolute $EC_{50}/IC_{50}$ | Effective – Random | 2.21e-58 | 7.42e-58 | 5.14e-72 |
| | Effective – Border | 0.00e+00 | 0.00e+00 | 5.16e-206 |
| | Random – Border | 0.00e+00 | 0.00e+00 | 7.93e-143 |

Table 1: p values for the obtained relative $EC_{50}/IC_{50}$ and absolute $EC_{50}/IC_{50}$ for dose response curves with 8 doses, 1, 2, or 3 replicates, 20 negative controls on 384-well plate, and strong plate effects with a linear relationship to column number on the right side of the plate.

| | 1% | 5% | 10% | 40% |
|---|---|---|---|---|
| Border | 0.92 ± (0.014) | 0.92 ± (0.007) | 0.92 ± (0.008) | 0.92 ± (0.004) |
| Random | 0.95 ± (0.010) | 0.95 ± (0.005) | 0.95 ± (0.004) | 0.94 ± (0.002) |
| Effective | 0.98 ± (0.007) | 0.98 ± (0.002) | 0.97 ± (0.002) | 0.98 ± (0.001) |

Table 2: Analysis of the AUC obtained for 10 screening experiments with varying hit rates, using layouts with 10 positive and 10 negative controls in the presence of very strong bowl-shaped plate effects. Each screening contains 40 plates. Hits are randomly located on the plates.

| | 1% | 5% | 10% | 40% |
|---|---|---|---|---|
| Effective – Random | $1.99 \times 10^{-06}$ | $7.78 \times 10^{-10}$ | $5.99 \times 10^{-11}$ | $1.75 \times 10^{-16}$ |
| Effective – Border | $1.56 \times 10^{-08}$ | $1.29 \times 10^{-10}$ | $1.38 \times 10^{-9}$ | $2.62 \times 10^{-12}$ |
| Random – Border | $4.64 \times 10^{-05}$ | $5.51 \times 10^{-8}$ | $4.04 \times 10^{-7}$ | $9.70 \times 10^{-10}$ |

Table 3: Analysis of the AUC obtained for 10 screening experiments with varying hit rates, using layouts with 10 positive and 10 negative controls in the presence of very strong bowl-shaped plate effects. Each screening contains 40 plates. Hits are randomly located on the plates.
